# A topic modelling analysis of TCGA breast and lung cancer transcriptomic data

**DOI:** 10.1101/2020.10.19.345694

**Authors:** Filippo Valle, Matteo Osella, Michele Caselle

**Affiliations:** Physics Department, University of Turin and INFN, via P. Giuria 1, 10125 Turin, Italy

**Keywords:** network-based cancer data analysis, topic modelling, gene expression, network theory, stochastic block modelling

## Abstract

Topic modelling is a widely used technique to extract relevant information from large arrays of data. The problem of finding a topic structure in a dataset was recently recognized to be analogous to the community detection problem in network theory. Leveraging on this analogy, a new class of topic modelling strategies has been introduced to overcome some of the limitations of classical methods. This paper applies these recent ideas to TCGA transcriptomic data on breast and lung cancer. The established cancer subtype organization is well reconstructed in the inferred latent topic structure. Moreover, we identify specific topics that are enriched in genes known to play a role in the corresponding disease and are strongly related to the survival probability of patients. Finally, we show that a simple neural network classifier operating in the low dimensional topic space is able to predict with high accuracy the cancer subtype of a test expression sample.

## 1. Introduction

Thanks to the impressive progress in sequencing technologies and the development of dedicated gene expression databases like *TCGA* [1], it is now possible to study in an unified way the transcriptional status of thousands of cancer samples for several different biological conditions and cancer types. These transcriptomes provide a huge amount of information to link pathological phenotypes to their molecular underpinnings and can be used to identify and classify cancer types and subtypes, find new biomarkers and, as a final goal, elaborate new therapeutic strategies. The early identification of the particular cancer subtype of a given patient may help to customize “ad hoc” therapeutic protocols and may greatly improve the survival probability of patients [2].

To address this issue one must deal with two main steps. First, one must identify the molecular signatures of cancer types and, possibly, of their subtype organization, by suitably clustering gene expression data. Second, one should use these signatures to train a classifier to allow a fast and reliable association of a new sample to the corresponding subtype. As an extra bonus, one may hope in this way to identify specific drivers of the particular cancer subtype under study and reconstruct a map of the altered pathways that promote tumor growth and aggressiveness. However, finding signatures able to distinguish among different cancer subtypes is a highly non trivial task. From a theoretical point of view it is a typical example of a dimensionality reduction process. Starting from the huge dimensional space of gene expression data (with thousands of genes) one aims to find a few (or orders of dozens) of relevant subsets able to summarize the whole information content of the original dataset.

In general a good signature is a collection of genes which are able to capture a large portion of the variation in gene expression across tumors of the same type. These genes can thus be used to define and identify molecular subtypes of that cancer and may explain (and predict) the clinical heterogeneity of the disease (for instance different levels of overall survival). There are several methods for selecting gene signatures. Most of them are based on unsupervised clustering algorithms [3], with a wide variety of different implementations.

Despite the apparent simplicity of this program, its actual realization is not so easy. It often happens that lists of genes obtained by different labs and putatively associated to the same cancer subtype show almost no overlap among them [4]. This is due to the high heterogeneity at the molecular level of tumors [5] and the intrinsic complexity of performing dimensionality reduction in the gene expression space.

In the past few years, a new class of clustering algorithms based on the so-called “topic modelling” approach has been proposed in order to address this issue [6]. This proposal stems form the observation that a similar degree of complexity and heterogeneity is also present in Natural Language Processing. Indeed algorithms which try to identify the “topics” associated to a given document from the word usage have to face the same type of challenges we are facing here. In this analogy the cancer samples play the role of the documents, the words are the genes, the number of times a particular word is used in a given document is the analogous of the expression level of a particular gene in a given sample. The topics are the gene sets (the “signatures”) we use to cluster samples into subtypes. The goal of topic modelling is to identify the “topic” of a given document from the word usage exactly as our goal is to identify the cancer subtype from the gene expression pattern. The major advantage of topic modelling methods with respect to standard clustering approaches is that they allow a “fuzzy” type of clustering [7]. The output of a topic modelling algorithm is a *probability distribution* of membership, i.e., the probability of a given document to be composed by a given set of topics and, at the same time, the probability of a word to characterize a given topic. In our context, this means that we have as output a set of values that quantify the probability of a given sample to belong to a particular cancer subtype and the relevance of a given gene in driving this identification.

The most popular tool to perform this kind of analysis is the so-called Latent Dirichlet Allocation (LDA) algorithm, which basically assumes a Dirichlet prior distribution for the topic distribution. There is no particular motivation in the natural language context as well as in our biological context for the Dirichlet prior. Its motivation comes from the fact that it allows a simple solution of the allocation problem and thus the algorithm can be efficiently applied to databases with a large number of documents and words. However, the lack of biological motivation for the prior and the large number of free parameters represent a possible limitation of LDA in our context.

Another common approach, often used in addressing gene expression data, is the Nonnegative Matrix Factorisation [8]. The major drawback of this approach is that it facilitates the detection of sharp boundaries among subtypes and this could be a limitation in very heterogeneous settings.

In recent years, some important advances in the field have laid the foundations for overcoming some of the limits of LDA. First, it has been realized [9,10] that there is a strong connection between topic modelling analysis of complex databases and the community detection problem in bipartite graphs, which is a well know and much studied problem in network theory [11]. Second, a very effective class of community detection algorithms, based on hierarchical stochastic block modelling (hSBM), has been adapted to the topic modelling task [9].

A major advantage of hSBM type algorithms is that they do not require any particular assumption on the probability distribution of the latent variables and can thus adapt to the possible heterogeneous nature of gene expression data in cancer cell lines. Moreover, they do not require external inputs such as the expected number of clusters (or topics) as it is the case for LDA and in general for standard clustering algorithms. They are able to recognize the hierarchical organization of samples (and genes) within the database and output an optimal choice for the number of topics at different levels of resolution (i.e., the algorithm is able to identify the hierarchical organization of the cancer samples in the database). The major drawback of this class of algorithms is the large amount of computing power required. Indeed, it was only in the last few years that it became feasible to address large and dense networks, like the ones in which we are interested, with hSBM methods.

In this paper we apply for the first time hSBM-based topic modelling to the study of cancer gene expression data. This paper is part of an ongoing effort in our lab to explore the possible applications of advanced network based computational tools to the molecular characterization of cancer and, in particular, to the identification of cancer drivers [12–15].

This paper focuses in particular on breast and lung cancer, due to their clinical relevance and to the presence of a large number of studies which can be used to benchmark our result. However, the methods developed here could be in principle applied to any type of cancer.

Our main goal is to identify signatures for breast and lung cancer subtypes and then use these signatures to construct an efficient classifier to associate samples to their most probable subtype. We shall reach this goal by studying the gene content of the topics associated to the various subtypes. The analysis of the resulting “signatures” bring information on the peculiar features of the gene expression patterns in specific cancer cell lines.

By comparing our results with those of standard clustering methods we shall show that hSBM can infer information, like the optimal number of clusters, that cannot be obtained with other approaches. Finally, we will investigate how topics and clusters are related to the survival probability of patients.

## 2. Results

### 2.1. A topic modelling analysis of TCGA gene expression data of breast and lung

It has been recently realized [9] that there is a strong connection between topic modelling analysis of complex databases and the community detection problem in bipartite graphs. Among the several different approaches to community detection [11] one of the most powerful is the so-called hierarchical Stochastic Block Model (hSBM) [16], which belongs to the class of probabilistic inference approaches to community detection. This method has the advantage of making the minimal amount of assumptions on the data structure. A suitable implementation of this algorithm on bipartite networks leads to a particularly effective version of topic modelling that is able to identify at the same time the hierarchical structure of the network and keep track of its bipartite nature [9]. As a consequence, it automatically detects the number of topics and hierarchically clusters both sides of the network (in our case both genes and samples), thus overcoming the typical limitations of standard topic modelling approaches.

These properties are perfectly suited to deal with gene expression because of the heterogeneous nature of these data and the inherent hierarchical structure (e.g., the organization in tissues or cell lines).

Moreover, they are particularly relevant if one is interested in cancer gene expression data for which this hierarchical structure also shows up in the cancer subtypes organization. In the framework of precision medicine, it is of utmost importance to be able to identify cancer subtypes in a reliable and reproducible way to improve therapeutic treatment and predict survival probabilities.

We address the problem of cancer subtype identification using a hSBM based topic modelling analysis of RNA-Sequencing samples from The Cancer Genome Atlas (TCGA). In particular we focused on the TCGA-BRCA, TGCA-LUAD and TCGA-LUSC projects. These data sets can be represented as bipartite networks relating genes *{g*_*i*_*}* with samples *{s*_*j*_*}*. Each link of the network has a weight *w*_*ij*_ encoding the expression level of the gene *g*_*i*_ in the sample *s*_*j*_.

The output of hSBM is a hierarchical and probabilistic organization of the data in “blocks”. Each sample and each gene has in output a certain probability to belong to a given block (see the description of hSBM in the Methods section). We define as *clusters* the blocks of samples and as *topics* the blocks of genes (see Figure 1). Genes can be associated to topics and samples to clusters with a hard membership, i.e., each gene is linked only to a given topic and each sample only to a given cluster. However, the algorithm can be naturally extended to a “fuzzy” version of membership in which, for example, a sample has a non-zero probability of belonging to different clusters. While this option would certainly be interesting in our context, it adds a further layer of complexity that we decided to postpone to a forthcoming study.

**Figure 1.**
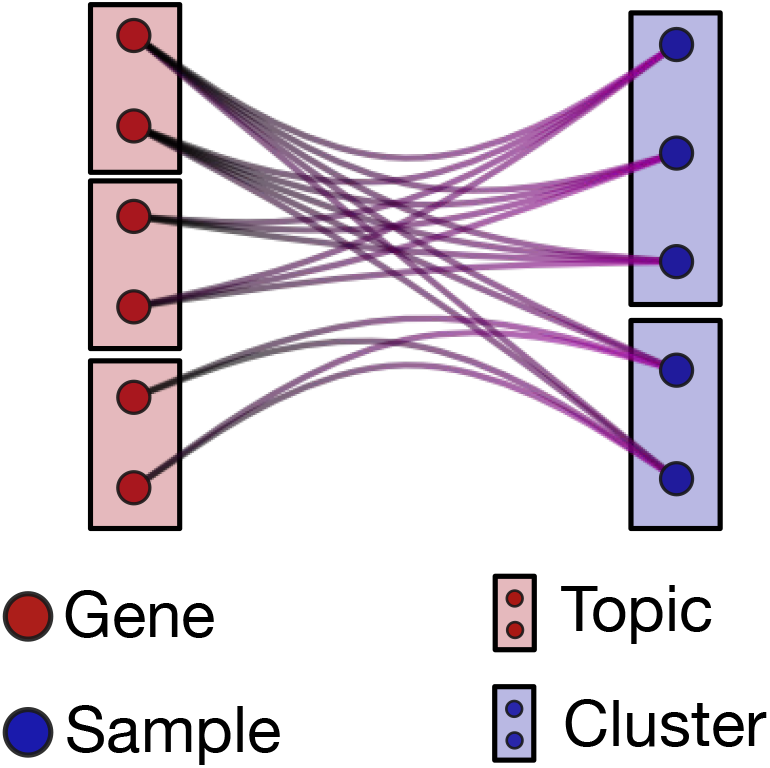
The hSBM partition of samples in clusters and genes in topics. The lines connecting genes and samples encode the weights of the bipartite network (i.e., the gene expression values in the different samples).

Therefore, the whole complexity of the database is finally encoded in the probability distributions *P*(topic|sample) and *P*(gene|topic) (see the Methods section for a precise definition).

- **P**(**topic**|**sample**) is the probability that a sample (or more precisely its overall gene expression pattern) is driven by the cooperative action of a particular set of genes (topic). These probabilities are not characterized by a hard membership but describe instead a “many to many” interaction (and for this reason they are able to capture the whole complexity of the problem). A given sample can be controlled by several different sets of genes (or topics) and a given set of genes can characterize different samples, possibly in different clusters, which in our context will identify different cancer subtypes. To better understand this result it may be useful to address the analogy with the standard topic modelling analysis of linguistic corpora. As mentioned in the introduction in this comparison genes are mapped to words and samples to documents, then *P*(topic|sample) gives us the probability that a particular document (sample) deals with a given topic. Similarly in our case this probability tells us the relevance of a set of genes (topic) in describing the sample and, as a consequence, in driving its assignment to a particular phenotype (cluster).
- **P**(**gene**|**topic**) is the probability distribution of the different genes within a topic. We mentioned above that genes are organized within topics with a hard membership partition. However the genes within a topic are not on the same ground, since their assignment to the topic is weighted by the *P*(gene|topic) distribution. This allows us to identify the genes (the ones with a larger *P*(gene|topic)) which play the most important role within the topic and are thus likely the drivers of the gene set effect on the samples. Indeed, we will show that by ranking the genes within a topic according to their assignment probability, the top ranking entries are well known oncogenes. Following again the above analogy with linguistic, *P*(gene|topic) is the analogous of the importance of a given word to define a topic. By selecting the top ranking words we may immediately understand the nature of the topic to which they are associated.

We shall concentrate in the following in particular on two cancer types: breast and lung (more precisely on the Non-Small-Cell Lung Cancer, which represents the majority of lung cancers). This choice is motivated by their clinical importance but also by the different and complementary challenges these cases present to our approach. While the subtype organization of breast cancer is based on gene signatures, the one of lung cancer is of clinical nature. While for the Non-Small-Cell Lung Cancer we only have two subtypes, in breast five different subtypes have to be identified, which represents a much more complex task.

We shall address in the following these two cases separately. We shall first evaluate the ability of hSBM to identify cancer subtypes and then we shall see what we can learn on the pathology (in terms of functional characterization of driving topics, new samples classification and survival prediction) from our analysis.

### 2.2. Gene selection

An important preliminary step of our analysis is gene selection. This reduction of the number of features is often a mandatory step due to the high computational cost of several algorithms. Among the various possible strategies, we here considered two alternative choices that are frequently used in this type of analysis: tissue specific genes or highly variable genes (see the Methods section for a precise definition and a detailed discussion). As we will show, a nice feature of hSBM is the robustness of its performances to the gene selection procedure. This should not be so surprising. First of all, the gene lists selected with the two criteria have often a substantial overlap. More importantly, a main message of our analysis is that the whole gene expression profile, and not only the behaviour of a small set of genes (e.g., markers) leads to the correct sample classification. Accordingly, the gene selection should not be a crucial step as long as the gene selected are sufficiently representative of overall expression program. For the sake of brevity, we will discuss in the main text the tissue-specific gene selection for the breast cancer case and the highly variable gene selection for the lung cancer case.

A comparison with the results using the alternative choice is reported in the Supplementary Material (see Figure S4).

### 2.3. Analysis of breast cancer samples

Breast cancer is the most common malignancy in women and one of the three most common cancers worldwide [17–19]. It is also one of the few examples of a tumor for which there is a widely accepted subtype classification [20–22] based on gene expression that has a relevant therapeutic role and is instrumental for better clinical outcomes (in particular for HER2 subtypes). Breast cancer samples are usually divided in 5 different subtypes: *Luminal A, Luminal B, Triple-Negative/Basal, HER2* and *Normal-like*. On the clinical side, this classification is based on the levels of a handful of proteins whose presence in the biopsy is usually detected using immunohistochemistry (IHC) assays. In particular two hormone-receptors (estrogen-receptor (ER) and progesterone-receptor (PR)), HER2^1^ and Ki-67 ^2^.

Here is some information on these subtypes and their clinical outcomes.

- **Luminal A** breast cancer is hormone-receptor positive, HER2 negative, and has low levels of the protein Ki-67. Luminal A cancers are low grade, tend to grow slowly and have the best prognosis.
- **Luminal B** is very similar to Luminal A from the gene expression point of view. The main difference is that it can be either HER2 positive or HER2 negative, and is typically characterized by high levels of Ki-67. As a consequence Luminal B cancers generally grow slightly faster than Luminal A cancers and their prognosis is slightly worse.
- **Triple-negative/Basal** (which we shall simply denote as Basal in the following) are both hormone-receptor negative and HER2 negative.
- **HER2** is hormone-receptor negative and HER2 positive. This class of breast cancers tend to grow faster than the Luminal ones and can have a worse prognosis, but they are often successfully treated with targeted therapies aimed at the HER2 protein.
- **Normal-like** breast cancer is similar to Luminal A: hormone-receptor positive, HER2 negative, and has low levels of the protein Ki-67. However, its prognosis is slightly worse than Luminal A prognosis.

The same classification can be obtained (to a large extent [23]) looking at the expression levels of the well known “Prediction Analysis of Microarray (PAM)50” signature [24,25]. Given the expression levels of these signature genes, samples are then classified using standard machine learning methods (Classification and Regression Trees (CART), Weighted Voting (WV), Support Vector Machine (SVM), Nonnegative Matrix Factorisation (NMF) or k-Nearest Neighbors (k-NN)) or using methods based on the euclidean distance in the signature space like Nearest Template Prediction (NTP) [26] or with more sophisticated network-based methods like Hope4genes [15]. The agreement among different classifiers and with the IHC based subtyping is in general reasonably good but far from perfect (see for instance [15] for a recent comparison of the performances of different classifiers in a set of breast cancer classifications tasks). Recently, a discrepancy between IHC subtypes and PAM50 intrinsic subtypes was examined in detail [27] further suggesting that the standard annotations should be probably revised in the future. The classification task is made particularly difficult by the heterogeneity of cancer tissues (biopsies may contain relevant portions of healthy tissue) and by the intrinsic variability of gene expression patterns in cancer cell lines. For instance, the TCGA samples that we shall use for our analysis have been recently reanalyzed in TCGABiolinks [28] leading to a significant relabeling of samples.

To address this particular issue, we downloaded both the *PAM50* labels from [29], which is the most widely used set of annotations, and the more recent and highly curated *SubtypeSelected* annotation provided by the new functionalities of TCGABiolinks [28]. In the following we shall compare the performance of our algorithms against both these annotations.

Our main goal in this framework is not to propose a new signature or a new classifier on top of the existing ones, but to show that it is possible to obtain relevant information on the cancer samples, like subtype annotation, the survival probability or lists of potential driver genes and altered pathways, **without** resorting to the marker genes mentioned above but looking instead at the **overall** gene expression pattern. We think this is an important achievement since it allows us to address breast cancer (and in principle any other complex pathology or cancer) without being influenced by the expression levels of few, often wildly fluctuating, marker genes, and opens the possibility to find new driver genes and possibly new subtype structures that may have therapeutic relevance.

We performed the hSBM analysis on a bipartite network starting from all the 1222 samples of the TCGA-BRCA project on one side and a suitable selection (see Methods section) of genes on the other side; the links were weighted by the expression values.

#### 2.3.1. Clustering of breast cancer samples

We clustered the 1222 samples of the TCGA-BRCA project using hSBM and a set of other state of art clustering tools: Latent Dirichlet Allocation (LDA) [30], Weighted Gene Correlation Network Analysis (WGCNA) [31] and hierarchical clustering (hierarchical) [32]. We also compared the quality of clustering with the two annotations *PAM50* [29] and *SubtypeSelected* [28]. Supplementary Table S3 reports the number of samples annotated for each subtype in the two annotation systems.

On the gene side instead of looking to cancer specific markers we selected, as mentioned above, only breast related genes i.e., genes whose behaviour was different in breast tissues with respect to other tissues (see Methods section for a precise definition). Results with the complementary choice of highly variable genes can be found in the Supplementary Material. After this selection, we ended up with 978 genes. Among these genes only HER2 among the classic markers discussed above was present. However, it plays no special role in our analysis since its expression level is on the same footing of the other 977 genes. We stress again that our goal was to show that it is possible to identify relevant features of cancer samples without assuming previous knowledge on existing markers.

hSBM finds a first layer of clustering in which the samples are divided into 8 clusters and the genes are organized in 6 topics. It identifies a further more refined level of organization composed by 29 clusters and 41 topics ^3^.

Results are reported in Figure 2. Figure 2a and 2b report the subtype organization in clusters for the first two layers (8 clusters for Figure 2a and 29 clusters for Figure 2b). We see looking at these figures that hSBM is able to identify rather well Basal, HER2 and Normal-like samples, while it mixes Luminal A and B samples. For this reason we shall treat them as a single subtype “Luminal” in the following.

**Figure 2.**
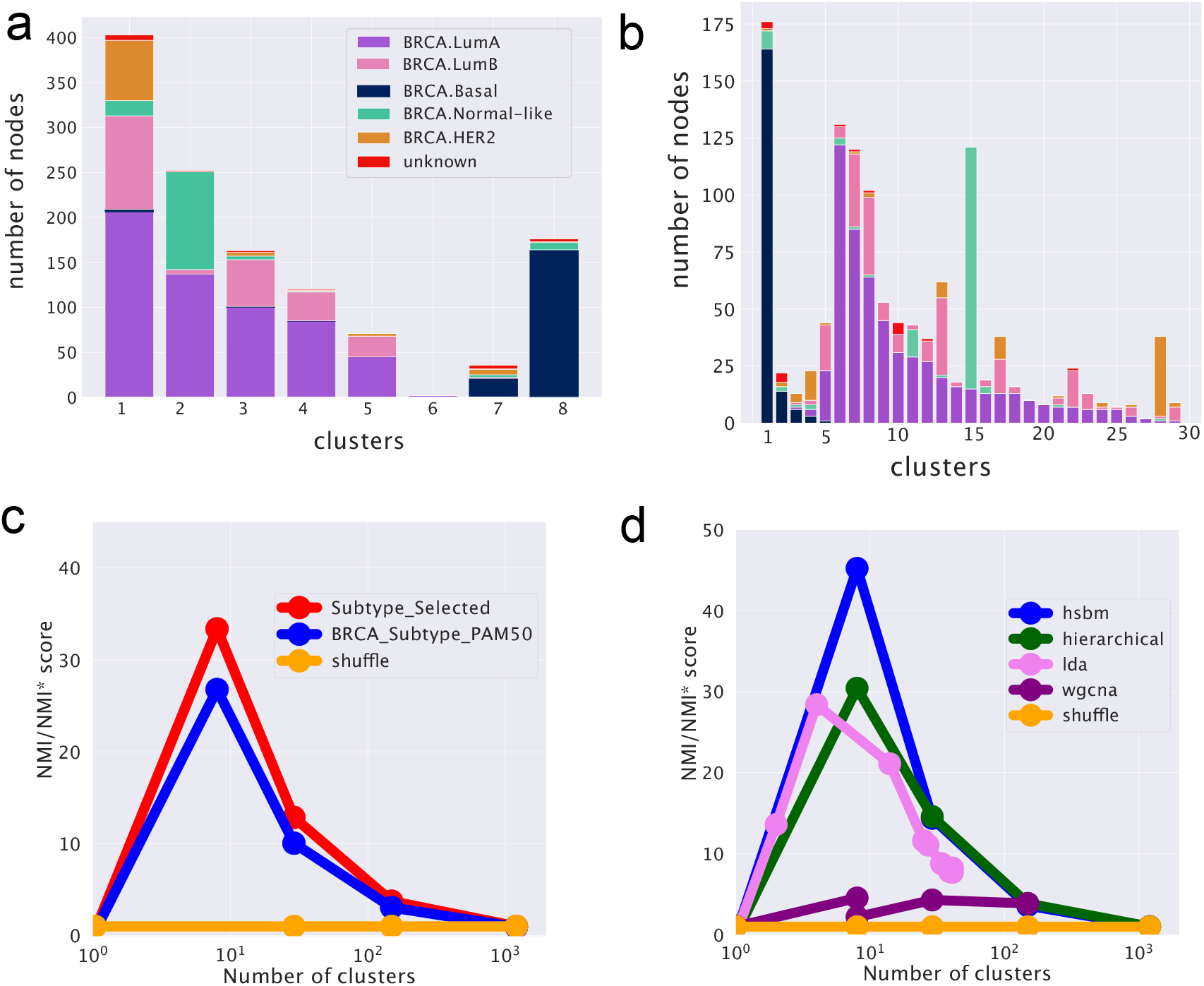
hSBM result for breast cancer analysis. In (**a**) and (**b**) it is reported the subtype composition of the clusters obtained via hSBM. Each column is a cluster, each color is a *SubtypeSelected* label from TCGABiolinks. The height of each column is proportional to the number of samples within the cluster. In (**a**) we report the results for the first layer of clustering (8 clusters) and in (**b**) those for the second layer (29 clusters). (**c**) Comparison of scores across hierarchy between TCGABiolinks *SubtypeSelected* labels [28] and TCGA labels from [29]. (**d**) Comparison of scores for different clustering algorithms. In (**c**) and (**d**) the NMI is scaled to the score obtained with a null model (NMI^*∗*^). See Methods section for more details.

We used the Normalised Mutual Information (NMI) measured compared to a null model as a score to evaluate the performance of the various algorithms in identifying cancer subtypes. NMI (see the Methods section) is a powerful tool to compare the performances of clustering algorithms in describing labelled partitions and was recently used to evaluate the performances of different topic modelling algorithms on synthetic corpora [33]. The NMI scores are reported in Figure 2d for the comparison between different clustering algorithms and in Figure 2c for the comparison of hSBM results using the two different sample annotations.

Looking at the figures we see that the highest NMI is reached for the first layer and that hSBM outperforms Weighted Gene Correlation Network Analysis (WGCNA), Latent Dirichlet Allocation (LDA) and hierarchical clustering. In order to set a comparison between different algorithms, the values of their several free parameters have to be selected. We chose the configuration of WGCNA, LDA and hierarchical clustering that could match more closely the number of topics and clusters obtained with hSBM. The rationale is to compare the different methods at the same resolution level (i.e., number of clusters and/or topics), thus at similar levels of dimensionality reduction. Therefore, it is possible that the algorithms could achieve better performances at different resolutions or using different performance metrics. This is for example the case of WGCNA. Setting WGCNA with different correlation threshold can improve its score but at the cost of producing in output a much larger number of topics and clusters with respect to hSBM. The Methods section and the Supplementary Figure S5 discuss more in detail this aspect.

Interestingly, we find a higher value of NMI at both resolution levels for the recent and more refined annotation of samples *SubtypeSelected* [28] with respect to the old one from [29]. In [28], the authors recognised several additional Normal-like samples thanks to an extensive effort to systematically quantify tumor purity with a variety of methods integrated into a consensus approach across TCGA cancer types. Indeed, the tumor microenvironment includes non-cancerous cells of which a large proportion are immune cells or cells that support blood vessels and other Normal-like cell. The Normal-like transcripts actually improve the results on the cancer classification.

Let us stress again that we obtained these results **without** resorting to the marker genes HER2, KI-67 and hormone receptors, which, except for HER2, were actually excluded from the gene set used in the analysis. Our results show that the expression levels of these genes are crucial to distinguish between the two Luminal types, but they are not mandatory to identify the remaining subtypes and separate them from Luminal A/B.

#### 2.3.2. Gene expression pattern in the topic space and subtype specific topics

One of the advantages of using a topic modelling approach is that, as we mentioned above, each sample is a mixtures of topics described by the probability distribution *P*(topic|sample). It is easy to obtain the probability *P*(topic|subtype) by averaging *P*(topic|sample) over all samples belonging to the same subtype. By subtracting to *P*(topic|subtype) the mean value over the whole dataset, we find a new set of quantities, that we define as “centered” distributions *P*(topic|subtype) (see eq.4). This centered distributions allow to identify subtype specific topics as topics that are particularly enriched in samples belonging to a particular subtype, and thus potentially playing a role in driving the specific features of the subtype. We report as an example in Figure 3 the results of this analysis for one particular topic (labelled as “Topic 1”) out of the 41 that we obtained at the second hierarchical level of our analysis. This topic turns out to be particularly enriched in the Normal-like subtype and slightly enriched in the Luminal A subtype. Other examples are reported in Figure 4.

**Figure 3.**
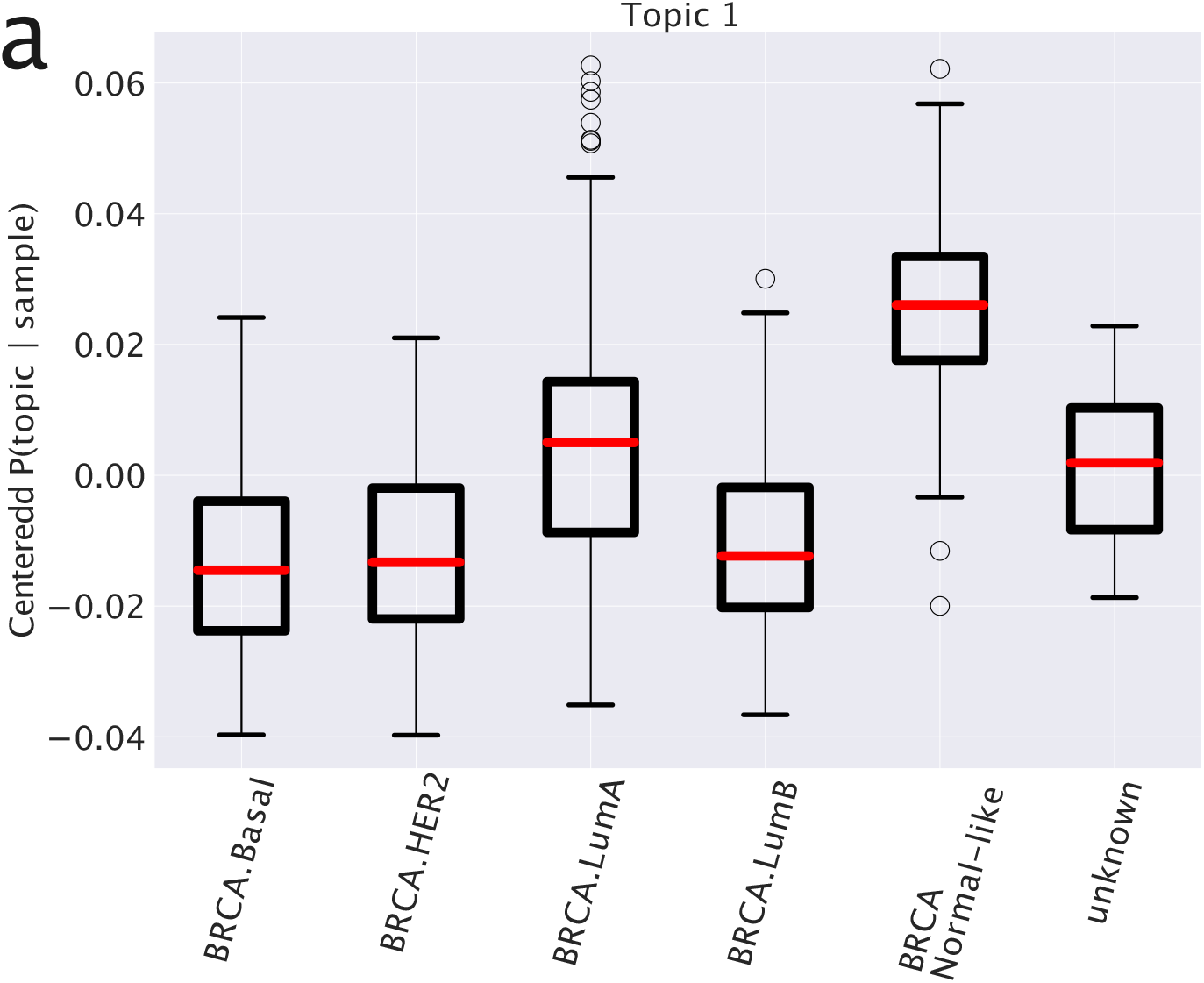
Values of 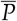 (topic|subtype) for Topic 1 and the different subtypes (see the main text for a detailed explanation). This centered version of the probability distribution allows to recognise the differences of topic expression in different subtypes.

**Figure 4.**
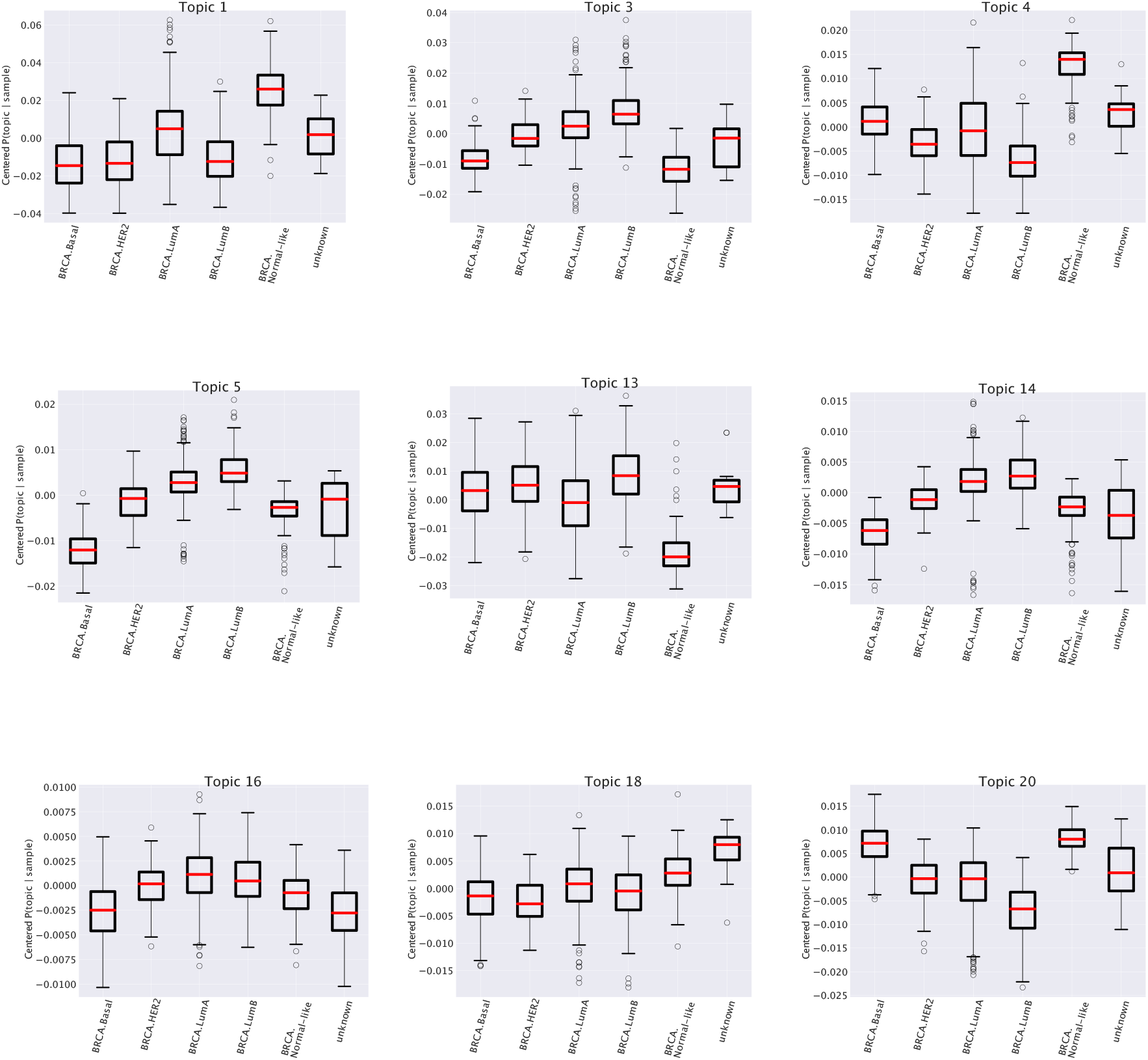
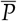(topic|sample) for various topics at the second level of resolution of hSBM (the same of Figure 3). Looking at the box plots we recognise several non trivial relationships between topics and subtypes. These relationships are consistent with the functional enrichment listed in Table 1.

#### 2.3.3. Functional enrichment of the topics

Topics are nothing but lists of genes. A common way to investigate their properties is to perform enrichment tests using tools like GSEA [34]. The enrichment analysis on genes associated to subtype-specific topics finds functional categories that are precisely in agreement with the specific annotations of the subtype. Some illustrative examples are reported in Table 1. For instance, the first entries of the table, corresponding to the Topic 1 mentioned above, show a strong enrichment for two sets of genes (labelled, following the GSEA convention as SMID_BREAST_CANCER_LUMINAL_A_UP and SMID_BREAST_CANCER_NORMAL_LIKE_UP) taken from [35] which fully agree with our subtype annotation in the topic space.

**Table 1.** Gene ontology results for few topics associated to a number of genes large enough for a reliable Gene Set Enrichment Analysis. We report only the entries with the most significant enrichment. In brackets the number of genes in each set (topic). Many topics are enriched for terms related to particular subtypes of breast cancer and this relation is indeed confirmed by the box plots in Figure 3 and 4. In some other cases, as for instance in Topic 13, entries such as *epithelial mesenchymal transition* are generic hallmarks of invasiveness or, possibly, of the metastatic nature of the sample and accordingly (see the box plot in Fig.4) they are shared by different subtypes.

| Term | FDR q-value |
| --- | --- |
| Topic 1 (72) |  |
| SMID_BREAST_CANCER_LUMINAL_A_UP | $3.03E - 40$ |
| SMID_BREAST_CANCER_NORMAL_LIKE_UP | $3.23E - 15$ |
| Topic 3 (50) |  |
| SMID_BREAST_CANCER_BASAL_DN | $4.89E - 9$ |
| VANTVEER_BREAST_CANCER_METASTASIS_UP | $3.46E - 8$ |
| Topic 4 (24) |  |
| SMID_BREAST_CANCER_NORMAL_LIKE_UP | $6.16E - 11$ |
| SMID_BREAST_CANCER_LUMINAL_B_DN | $1.34E - 8$ |
| Topic 5 (21) |  |
| VANTVEER_BREAST_CANCER_ESR1_UP | $3.03E - 40$ |
| SMID_BREAST_CANCER_BASAL_DN | $3.23E - 15$ |
| SMID_BREAST_CANCER_LUMINAL_B_UP | $1.41E - 6$ |
| Topic 13 (129) |  |
| HALLMARK_EPITHELIAL_MESENCHYMAL_TRANSITION | $7.8E - 14$ |
| SCHUETZ_BREAST_CANCER_DUCTAL_INVASIVE_VE_UP | $7.8E - 14$ |
| GO_EXTRACELLULAR_MATRIX | $6.9E - 12$ |
| Topic 14 (20) |  |
| SMID_BREAST_CANCER_BASAL_DN | $1.47E - 8$ |
| SMID_BREAST_CANCER_LUMINAL_B_UP | $3.04E - 6$ |
| VANTVEER_BREAST_CANCER_ESR1_UP | $5.66E - 4$ |
| Topic 16 (19) |  |
| MODULE_55 | $2.24E - 5$ |
| Topic 18 (42) |  |
| GO_CONTRACTILE_FIBER | $1.7E - 4$ |
| GO_SARCOPLASM | $3.56E - 4$ |
| Topic 20 (37) |  |
| SMID_BREAST_CANCER_LUMINAL_B_DN | $1.7E - 15$ |
| SMID_BREAST_CANCER_BASAL_UP | $3.3E - 10$ |

**Table 2.**
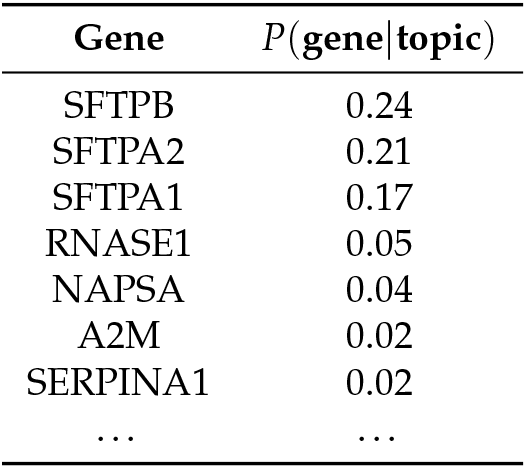
Genes in our candidate topic. Genes are sorted by their contribution to the topic. The complete list is available at https://github.com/fvalle1/topicTCGA/blob/master/lung/topic_3_lung.csv

This correspondence between subtype annotation of topics and functional enrichment of their gene content is further confirmed by the other entries of Table 1 and Figure 4. In particular we see, looking for instance at the last entry (“Topic 20”) of the table that it works also when the subtype is *depleted* in the topic space (see the last plot of Figure 4 in which the subtype Luminal B is depleted) to which corresponds an enrichment in *downregulated* genes in Luminal B breast cancer according to [35]. It is worth mentioning once again that in our analysis we selected only genes which are generically expressed in breast and not specifically differentially expressed in breast cancer. This makes the above results a non trivial consistency check of our procedure and further supports our idea of a role of the whole gene expression pattern of the cell in driving breast cancer subtype phenotypes.

#### 2.3.4. Predicting breast subtype annotation

One of the advantages of a topic model approach is that it is also a dimensionality reduction process. Topics can be interpreted as new coordinates one can use to visualise and study the data.

We used the topic space as an embedding space to train a neural network model which can then be used as an efficient classifier to associate samples to their specific subtype. Using topics as features and *SubtypeSelected* as labels our task becomes a simple supervised learning classification problem. The use of the topic space greatly simplifies the data space, and therefore the classifier can be trained much faster and with fewer parameters. Moreover, we showed that the topics have a non-trivial biological meaning and this can help the classifier in identifying the relevant structures in a possibly noisy data set. We obtained a high accuracy classifier using a 399 dimensional topic space (the lowest level of the hierarchical organization of the topic space) starting from a space with almost 20000 genes. Figure 5 reports in detail the performance of the classifier.

**Figure 5.**
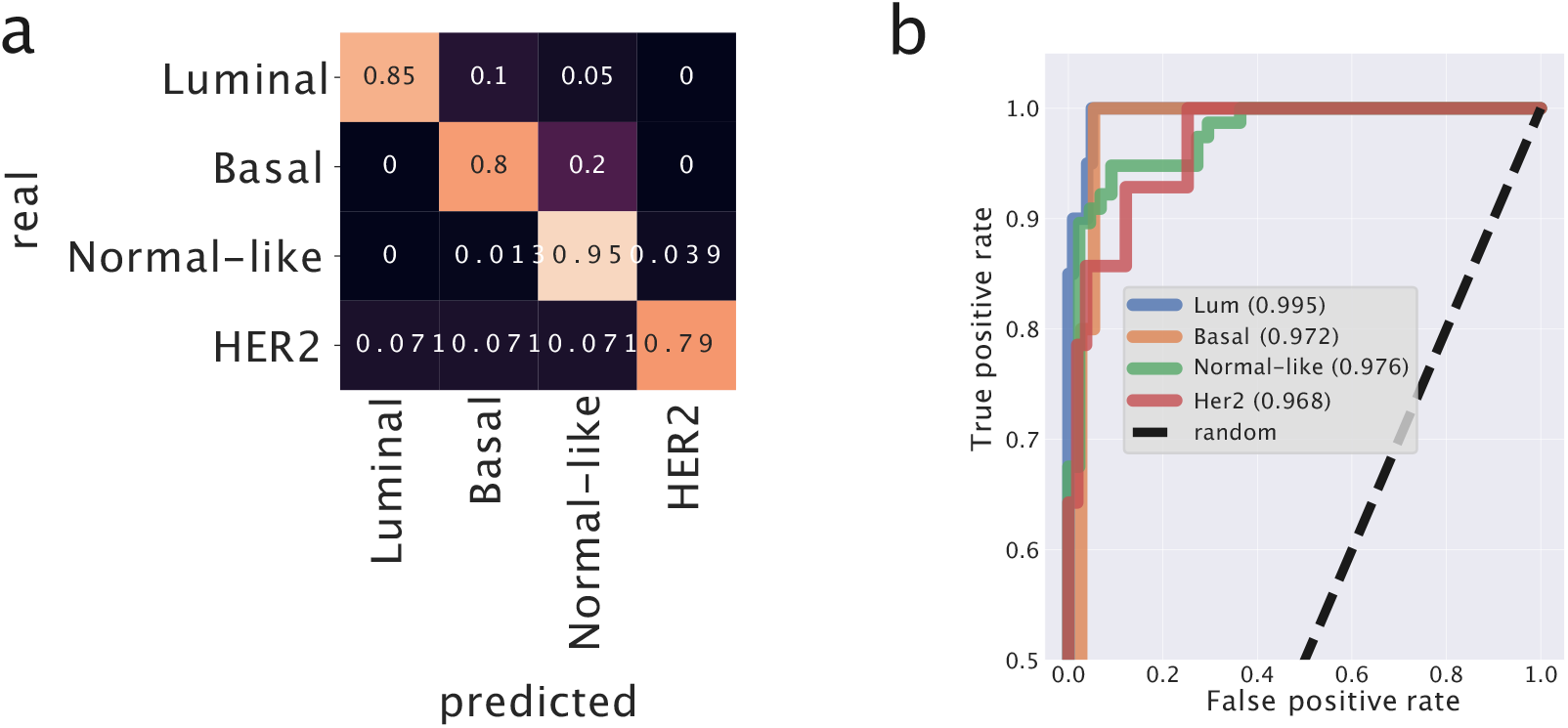
Predictor model for breast cancer. We built a neural network and trained it on the low-dimensional topic space to classify the different breast cancer subtypes. In (**a**) we report the confusion matrix. In (**b**) the Receiving Operation Characteristic curve and the corresponding Area Under Curve for each subtype estimated using a One-vs-All strategy four times. The diagonal (*TPR* = *FPR*) corresponding to random guessing is reported for reference. *Luminal* and *Basal* subtypes are the ones with the lowest fraction of False Positives. *Normal-like* is the subtype with the highest fraction of True Positives.

To evaluate the performances of our predictor we performed the same analysis using K-Nearest Neighbors which is a standard and very popular tool in this context. It turns out that the performances of our predictor model (AUC = 0.98) are greater than those of k-NN (AUC = 0.90) on the same dataset. This tells us that the organisation of the samples in the original gene expression space is not trivial, and that the projection into the topics space improves the ability of the predictor to assign the correct labels to test samples. We report the AUC score since it is less influenced by unbalanced classes than, for instance, the accuracy score. We applied K-NN using 5 `n_neighbors` and using *euclidean* `metric` on the log_2_(*FPKM* + 1) transformed data.

### 2.4. Analysis of Non-Small-Cell Lung Cancer samples

To reveal the potentialities of topic modelling in a different context, we analysed Non-Small-Cell Lung Cancer data taken again from TCGA. Lung cancer subtypes are currently defined by their pathological characteristics. The two predominant histological phenotypes of Non-Small-Cell Lung Cancer are adenocarcinoma and squamous cell carcinoma [36]. TCGA-LUAD and TCGA-LUSC projects provide transcripts for samples of these two subtypes. In the same way as in the breast analyses we collected FPKM data with GDC’s tools. In this case we selected 3000 highly variable genes (the second of the two options mentioned in the introduction). Results with the other choice, a tissue specific selection of genes, can be found in the Supplementary Material in Figure S4.

The binary choice (LUAD versus LUSC) represents a much easier task for a clustering algorithm and indeed, as we shall see, hSBM is able to correctly separate LUAD from LUSC. The TCGA repository on lung cancer data allows for a non-trivial test of clustering algorithms. Indeed, Cline et al. in [37] observed that some samples from TCGA-LUSC have gene expression levels that are more similar to LUAD than LUSC, although their similarity to LUAD is modest. On this basis they suggested that these samples may be borderline for subtype classification, for example because representing tumors that are less differentiated and thus difficult to classify by pathology. The list of these samples, labelled as *Discordant LUSC*, is provided [37]. We analysed how hSBM actually classifies these samples. Finally, in the context of lung cancer, thanks to the recent work of [38], we may perform another non trivial test of our topic modelling approach. We can combine together healthy and cancer tissues and look at the ability of hSBM to separate healthy samples from cancer ones.

#### 2.4.1. Classification of Non-Small-Cell Lung Cancer subtypes: LUAD and LUSC

Running hSBM on the above data we found three different layers of resolution with 2, 8 and 58 clusters and 5, 12 and 42 topics respectively.

The results of our analysis are reported in Figure 6 and 7. In particular, Figure 6a shows that hSBM is able to separate well the two subtypes and that indeed most of the *Discordant LUSC* samples are clustered together with the LUAD ones, capturing the fact that they are more similar to LUAD than LUSC. It is instructive to follow the hierarchical organization of clusters (Figure 6b). Already in the first layer many LUAD samples are separated from LUSC, while in the next layers the separation is complete.

**Figure 6.**
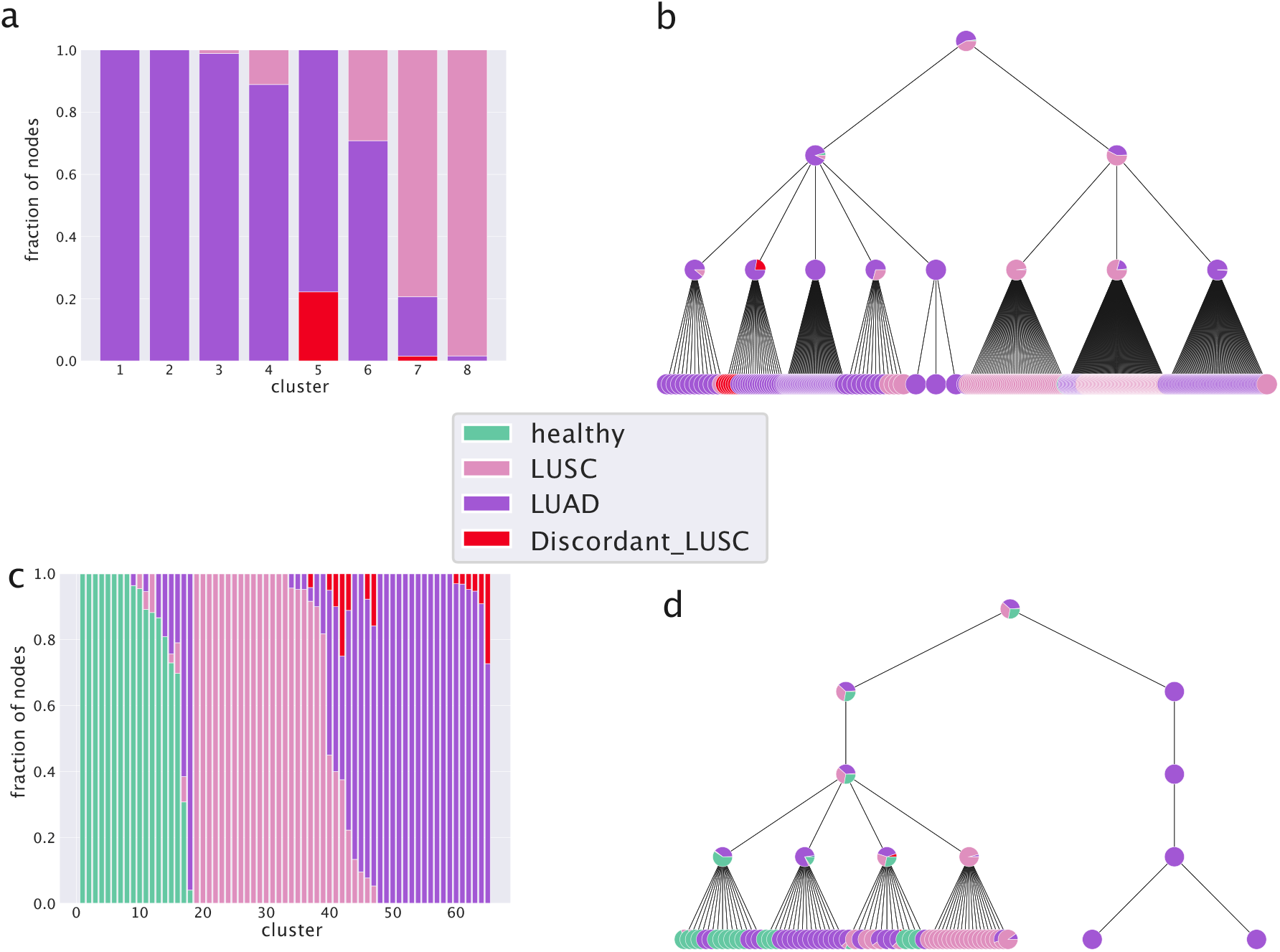
The hSBM classification of Non-Small-Cell Lung Cancer subtypes. In (**a**) the columns represent clusters and the colours refer to the sample annotations. Columns are normalized to the total number of samples in the cluster so that the height of different portions of the column are proportional to the fraction of LUAD or LUSC samples in that cluster. (**b**) The hierarchical structure. Already in the first layer many LUAD samples are separated from LUSC, in the next layers the separation is complete. In (**c**) and (**d**) we report the results of hSBM analysis including also healthy samples. In both settings *Discordant LUSC* are classified with LUAD.

**Figure 7.**
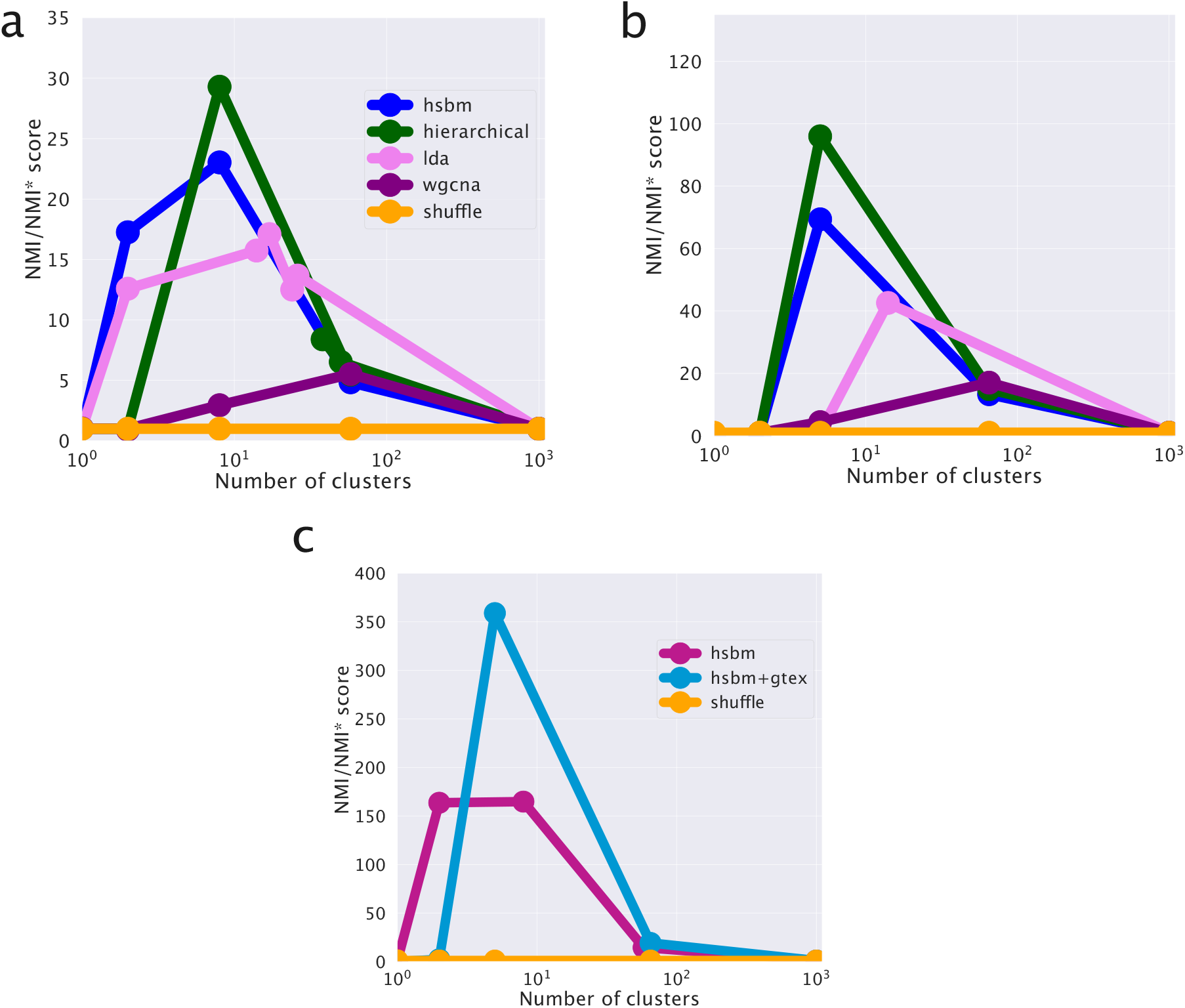
Comparison of different clustering algorithms. In (**a**) we report the scores for the classification of Non-Small-Cell Lung Cancer subtypes without healthy tissues. In (**b**) the scores in presence of healthy tissues. (**c**) The direct comparison between the case with and without healthy samples shows that their addition improves the cancer classification. Note that the score on the y-axis is normalized with respect to a case-dependent sample reshuffling (as explained in detail in the Methods section). This explains the different ranges of the scores in the panels.

As we did in the breast cancer case, we can study the distribution of subtype probabilities across topics and their enrichment in LUAD and LUSC samples. Two examples of topics that look differentially represented in the subtypes are reported in Figure 8.

**Figure 8.**
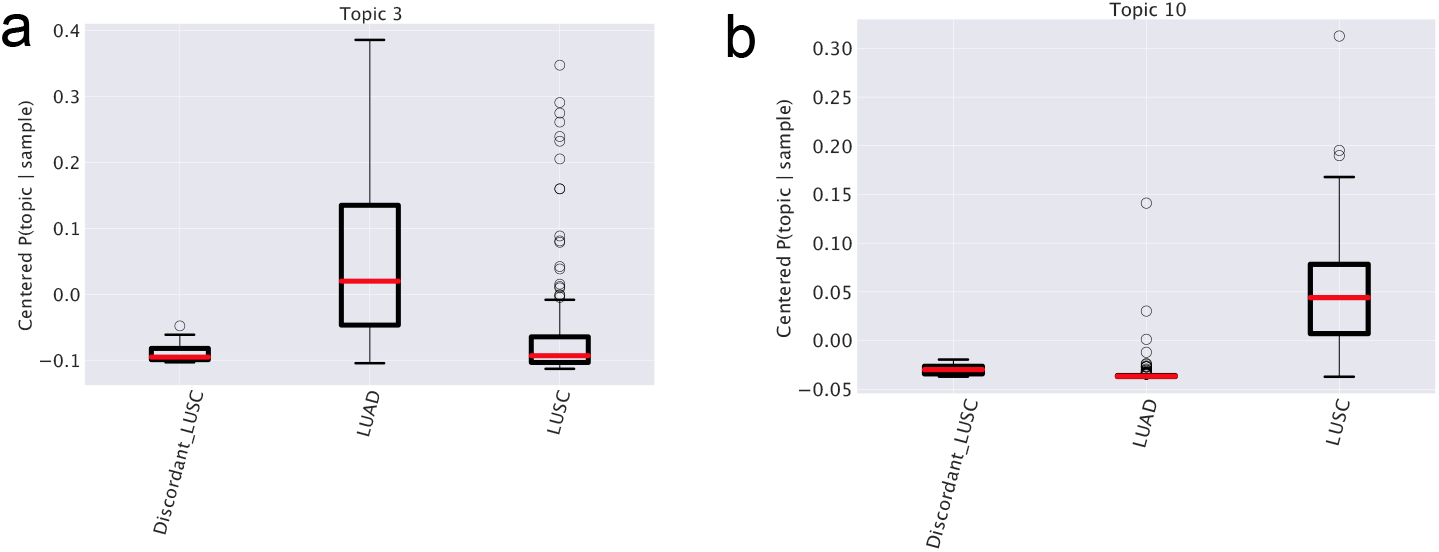
Topic trends in lung subtypes. In (**a**) and (**b**) the values of 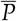 (topic|subtype) are reported for two topics grouped by the different subtypes (LUAD, LUSC and Discordant LUSC). The topic enrichment in different subtypes emerges.

#### 2.4.2. Classification of LUAD and LUSC samples versus healthy tissue

We also tested the ability of clustering algorithms and in particular hSBM to separate healthy from cancer tissues. We downloaded data with healthy (taken from the Genotype Tissue Expression [39] GTEx project) and cancers samples provided in a unified framework by [38,40]. We selected only samples with valid metadata available from TCGABiolinks or GTEx. Looking at Figure 6c we see that also in this case hSBM identifies LUAD and LUSC subtypes and that both are separated from healthy samples. The majority of *Discordant LUSC* samples are clustered with LUAD as discussed before. Even if in principle the task of identifying three categories instead of two is harder, it seems that the inclusion of healthy tissues actually improves the performance of hSBM. Looking at Figure 7c we see that the scores are higher than without healthy tissues.

Figure 7 shows that in the lung setting hSBM outperforms both LDA and WGCNA and it is compatible with hierarchical clustering.

As discussed in the breast cancer analysis, WGCNA is set to match the hSBM automatically retrieved number of topics and clusters. Note that WGCNA with very relaxed thresholds on the correlations becomes essentially equivalent to hierarchical clustering as it is shown in the Supplementary Figure S5. However, the relatively high performance score on the sample cluster structure comes at the cost of predicting a much larger number of topics (90) with respect to the 5, 12 and 42 topics retrieved by hSBM in the three different layers of resolution. A similar warning also holds true for LDA. In order to make a fair comparison with hSBM which has no free parameter, we used LDA with parameters set to their default values. In principle, LDA performances could be improved by suitably fine tuning its parameters, but such a extensive parameter exploration was beyond the scope of the present paper.

#### 2.4.3. Predicting LUAD and LUSC

The topic embedding space can be used to build a predictor analogous to the one we developed for breast cancer. In this case, the goal is to classify correctly LUAD and LUSC.

This predictor is actually a neural network with one hidden layer composed by 20 neurons and an output layer activated by a sigmoid function for the binary classification. We report in Figure 9 the output of this model on the test set. LUAD and LUSC are classified with high accuracy (accuracy: 0.9268, AUC: 0.9493). Also in this case, we compared our results with a standard K-NN predictor. K-NN achieves slightly better performances (accuracy: 0.9756, AUC: 0.9733). This is probably due to the fact that this task, which involves only a binary choice, is relatively simpler than the one we studied in the breast cancer case.

**Figure 9.**
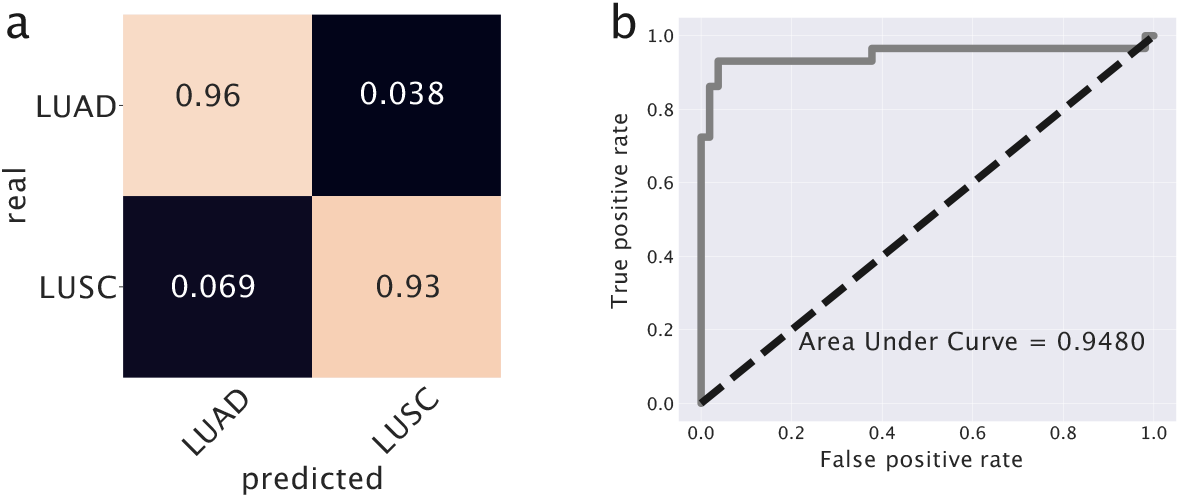
Prediction model for lung cancer. In (**a**) we used topics as features to train this model and we report the confusion matrix. In (**b**) the Receiving Operation Characteristic curve is reported. The Area Under Curve is reported as a score.

The classifier we are using is inherently probabilistic since in output gives the probability that a sample belongs to a specific class. When the difference between tumours subtypes is not so clearly defined, but there is instead a continuum of possible cancer types, a careful analysis of the actual classification probabilities can be informative. This is the case for the classification of the *Discordant LUSC* samples mentioned above. Figure 10 reports the algorithm output *Z*, which should be interpreted as the probability of a sample to be of LUSC type. The *Discordant LUSC* samples have a score in the range 0.3 *−* 0.4 and interestingly seem to emerge as an intermediate peak in the classification probability distribution. They typically have a probability to be LUSC greater than standard LUAD samples, even if this probability is below the classic 0.5 threshold for LUSC classification.

**Figure 10.**
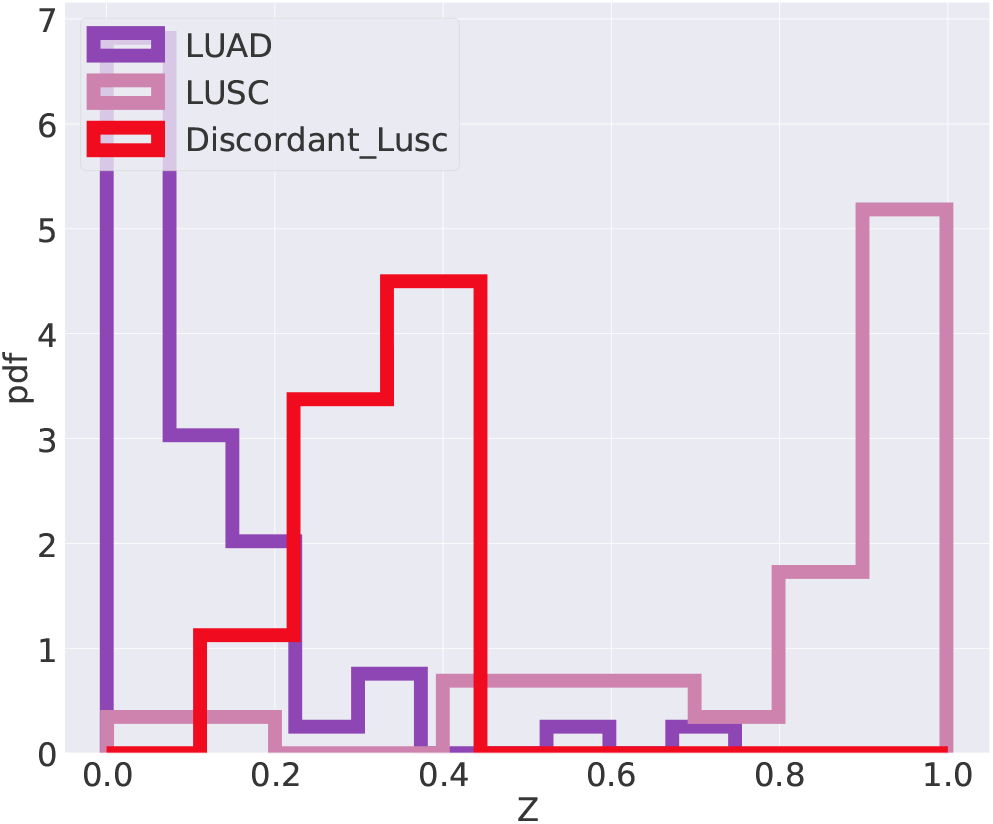
Output of the last layer of the predictor. *Z* is the output of the sigmoid function on the last layer *Z* = *σ*(*z*), namely *Z* is the probability of being LUSC.

## 3. Discussion

We saw that a topic modelling analysis is able to extract a lot of relevant information from cancer gene expression data. This information is encoded in the topic distribution and more precisely in the probability distributions *P*(topic|sample) and *P*(gene|topic). We have shown that by projecting the data in the topic space it is possible to build efficient predictors that can assign samples to the correct subtype. Moreover, by looking at the distribution of genes within the topics it is possible to extract relevant functional information on the subtypes.

We shall see now that if we include in the picture also some additional information on the stage of the tumor and on the follow up of the patients there are some further relevant clinical information that we can extract from the projection of the tumor samples into the topic space.

We shall see two examples, one for breast and one for lung cancer.

### 3.1. Survival analysis for breast cancer

Looking at Table 1 we see that one specific topic, namely “Topic 13”, is characterized by the typical annotations of invasive or already metastatic tumor. Figure 4 shows that this topic is almost equally distributed across all subtypes. However, such a condition should obviously have an important and pervasive effect in the transcriptome and hSBM should be able to detect this effect. Thus, we looked at the probability distribution of this topic on the clusters. We constructed the distributions *P*(topic|cluster) in analogy with what we did for *P*(topic|subtype). These distributions are plotted in Figure 11. This time we see a rather pronounced effect: a cluster (cluster 5) is relatively enriched in the topic, while the topic is strongly depleted in cluster 2 ^4^.

**Figure 11.**
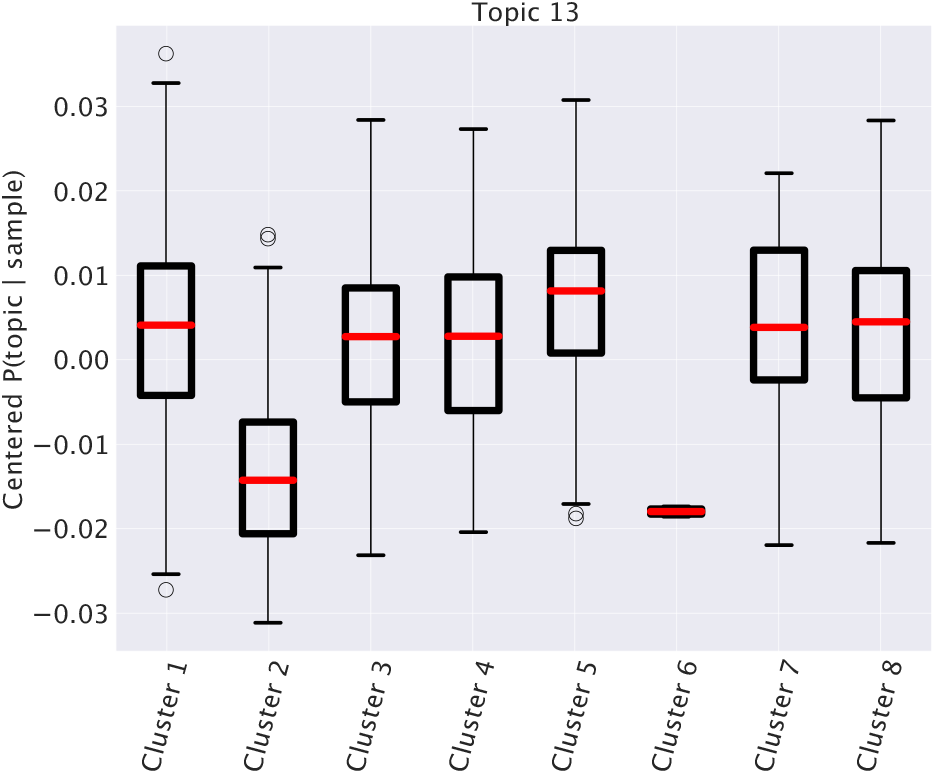
Topic 13’s expression in different clusters. 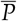 (topic 13|sample) for samples divided into clusters. The mixtures of samples of cluster 2 contains a lower fraction of topic 13 with respect to samples of other clusters.

At this point we can make a separate survival analysis for each of these clusters and the results are reported in Figure 12. We evaluated the significance of these results with a standard z-value reported in the figure. The score is obtained by running 1000 times a random reshuffling of samples annotations and keeping the size of clusters unchanged. The same analysis made at 3 years gives even higher z-values, in particular 3.13 for cluster 5 and 4.59 for cluster 2. Only these two clusters have a significant z-value. We see an impressive anticorrelation between the relevance of topic 13 in the cluster and the survival probability of the patients in the cluster. In particular, in cluster 2, a depletion of topic 13 was associated to a significant enhancement of survival probability.

**Figure 12.**
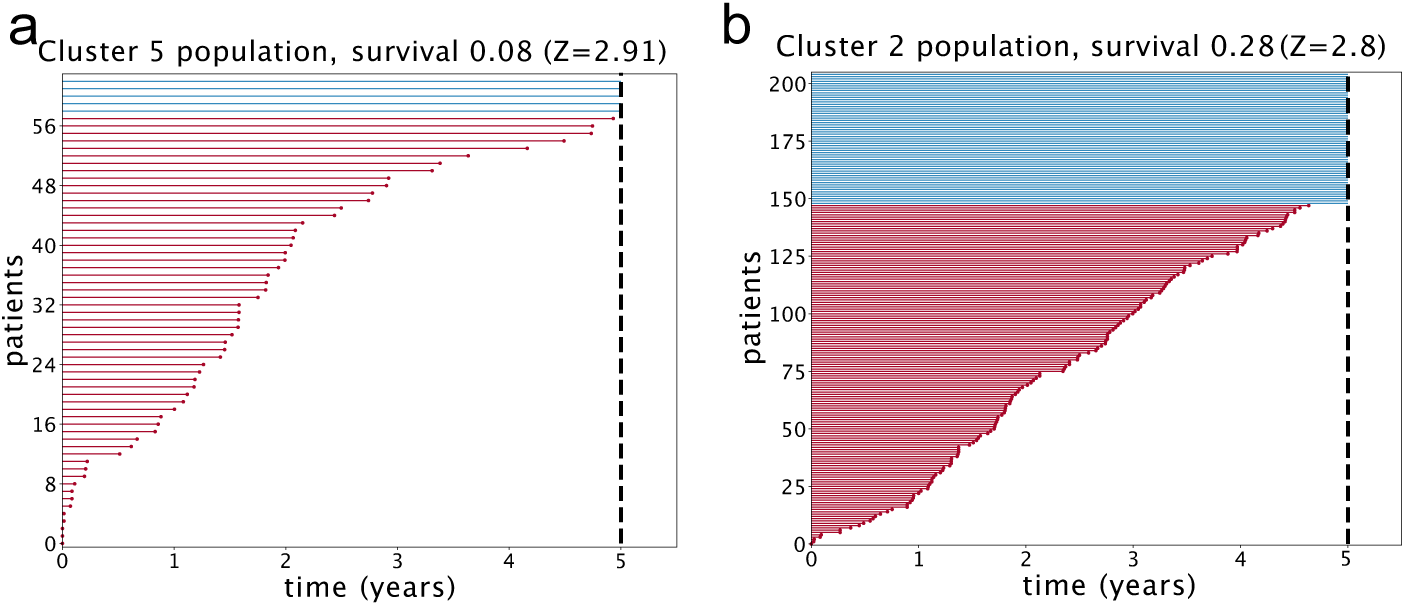
Survival time for two different clusters of patients. Each horizontal line is a patient; in red the ones who died or had the last follow up within 5 years from diagnosis, in blue the others. The survival is estimated as the fraction of patients that are alive after 5 years.

Interestingly, a similar analysis on the subtypes always gave less significant z-values in agreement with the above observation that Topic 13 is evenly distributed across subtypes. These analyses are reported in the Supplementary Figure S1.

We used the `plot_lifetimes` function of the lifelines package to obtain the plots in Figure 12. The *duration* argument was set to the number of days from diagnosis to death or to last follow up; the *event_observed* parameter was set to 1 for dead patients.

### 3.2. Survival analysis for LUNG cancer

Following a suggestion of Lucchetta et al. [41] we used the information on the tumor stage which is contained in the TCGA database to perform a survival analysis on the topic space.

Recently Lucchetta et al. [41] searched groups of genes effective in separating LUAD and LUSC. Their protocol consisted in extracting the genes that showed a trajectory of up-regulation across stages in one cancer type and down-regulation in the other.

Since topics are effectively group of genes we attempted a similar approach. We searched topics whose pattern across stages in the two different tumor subtypes was different.

In particular we focused on the topic (labelled as “topic 3”) in Figure 13, which shows, after an initial common trend, an up-regulation pattern through stages in LUAD and a down-regulation pattern in LUSC. We choose it as the candidate for the upcoming survival analysis. We provide in Supplementary Figure S2 the same analyses for all the other topics.

**Figure 13.**
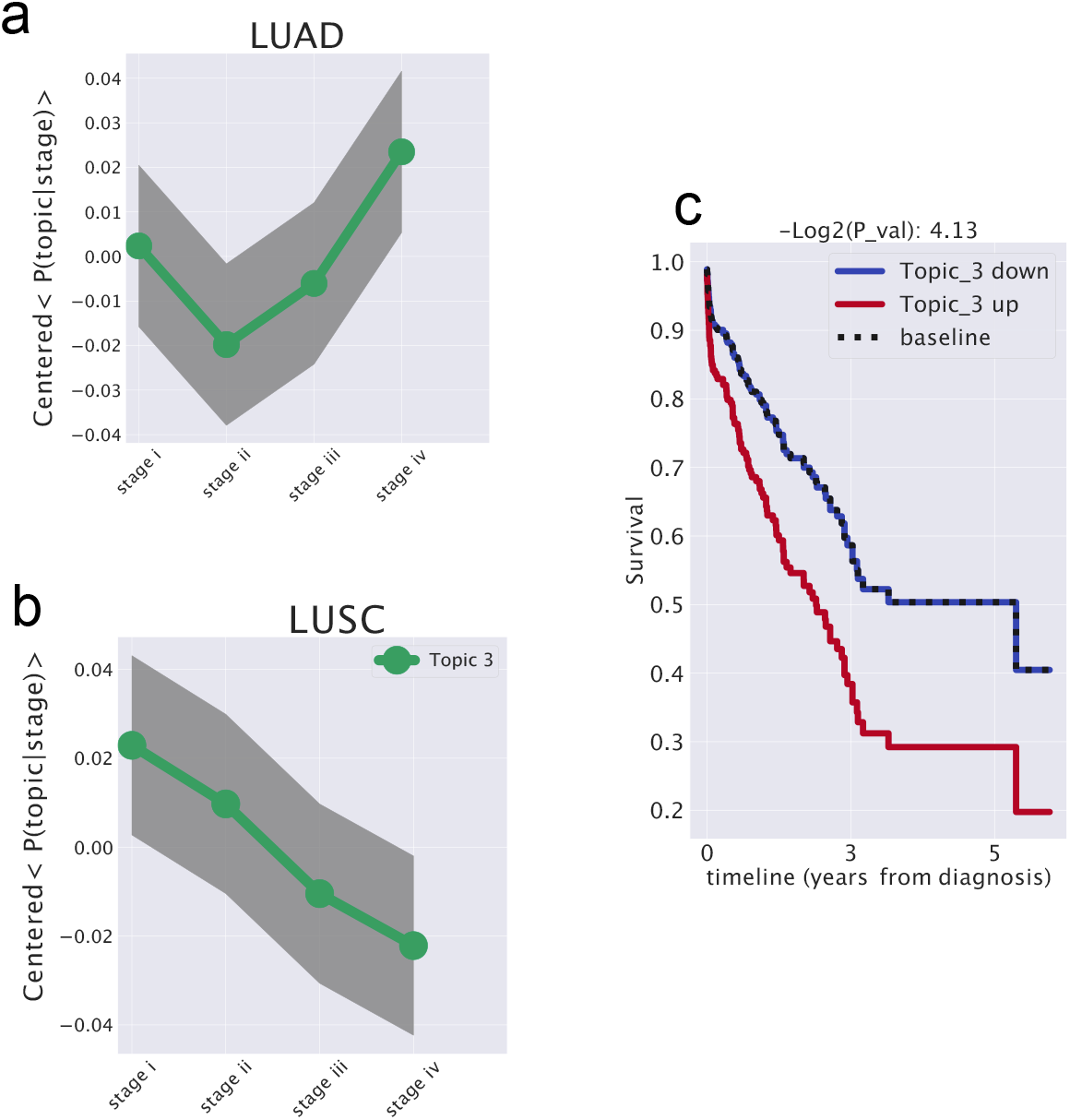
A candidate topic that affects patients’ survival probability and shows different trends in LUAD and LUSC. In (**a**) the trend of *P*(topic|stage) in LUAD. In (**b**) the trend of *P*(topic|stage) in LUSC. The genes belonging to this topic can be considered as a signature which distinguish the evolution across stages of the two subtypes. (**c**) The output of the survival analysis. When the topic is up-regulated the survival probability is halved.

The *lifelines* Python package was used to perform the survival analysis. We performed an estimation of the survival probability *S*(*t*), *cox* [42] was used to model the hazard function (the risk of dying at a certain time) and estimate how the survival probability is related to this topic. We defined a topic as up-regulated in a sample if the *P*(sample|topic) is above a threshold that corresponds to the 50th percentile. We found that patients with our candidate topic up-regulated had twice the probability to die with respect to the other patients. In order to test the significance of this results, we performed the same analysis using the *gender* variable instead of the topic up-regulation and found essentially no change in the survival probability between the two classes. As a further test we performed the same analysis for all the other topics at the same hierarchical level (see Figure S2 in the Supplementary Material). None of these other topics showed a different trend between LUAD and LUSC and accordingly we found no significant difference in survival in patients in which these topics were up-regulated (see Figure S2).

At this point we extracted the list of genes contained in topic 3 to see if we could understand the origin of this different survival probability. As we mentioned above, one of the advantages of topic modelling is that groups are not hard-constrained but are actually mixtures weighted by *P*(gene|topic): genes with the highest *P*(gene|topic) are likely to be those which contribute most to the topic.

We sorted the genes in this topic by *P*(gene|topic) and selected the first ones. Looking at the DISEASES database [43] we found that they are all related to lung cancer and lung diseases. Many terms like *cancer, bronchitis, pneumonia, interstitial lung disease* emerged.

Interestingly, the list of the genes contained in the topic has a very small overlap with the one found by Lucchetta et al. [41] even if both sets are able to successfully separate LUAD from LUSC and to predict different survival probabilities between the two. In particular, they analysed 2953 genes in 4 sets divided in up/down and LUAD/LUSC, we filtered 3000 genes; only 633 genes are in common, actually just 20% of genes are shared by the two analyses.

## 4. Materials and Methods

### Normalised Mutual Information (NMI)

We used the Normalised Mutual Information *NMI* [44] to evaluate the performance of different protocols to cluster together cancer subtypes. Given a set *C* of labelled samples and a partition *K* in clusters of these samples, the NMI is defined as the harmonic average of homogeneity *h* and completeness *𝒞*:

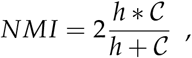

where the homogeneity is defined as

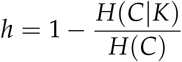

and the completeness as

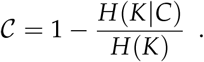

In these definitions *H*(*C*) and *H*(*K*) are the usual Shannon entropies associated to the partitions *C* and

*K*; *H*(*C*|*K*) and *H*(*K*|*C*) are defined as:

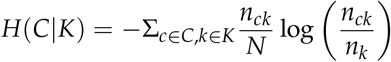

and

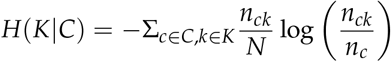

respectively, where *n*_*c*_ is the number of samples of the cancer subtype *c, n*_*k*_ the number of samples in the cluster *k* and *n*_*ck*_ the number of samples of the subtype *c* in the cluster *k*. With this definition it is easy to understand the meaning of homogeneity and completeness. If all the samples in cluster *k* belong to the cancer subtype *c* then *n*_*k*_ = *n*_*ck*_ and *h* = 1. Similarly the completeness *C* equals 1 if all samples belonging to the cancer subtype *c* are in the same cluster *k*. A similar approach involving *NMI* was proposed in [33] to evaluate topic modelling performance in reconstructing synthetic corpora.

Random clusters can have a non-zero NMI value given the way this score is defined. The score value will be a steadily increasing function of the number of clusters. To keep this effect into account, we normalized the empirical NMI values with the score NMI^*∗*^ obtained with a simple null model. The null model preserves the number of clusters and their sizes, but reshuffles the labels of samples. Thus, NMI/NMI^*∗*^ represents the how much the empirical score is higher than the score of random clusters of the same size and number. Moreover, we set the normalized score NMI/NMI^*∗*^ to 1 when both NMI and NMI^*∗*^ are zero, which is at the first layer where only one cluster is present. In order to perform a fair comparison of score values one should use the same null model for the normalization. Thus, for instance, the fact that adding GTEX data improves the performances of hSBM in the lung case cannot be deduced by simply comparing the absolute values of NMI/NMI^*∗*^ between Figures 7a and 7b. It is only the comparison in Figure 7c, performed with the same null model, which is fair and can be used to assess the improvement.

### TCGA data

The results published here are in part based upon data generated by The Cancer Genome Atlas (TCGA) managed by the NCI and NHGRI. Information about TCGA can be found at http://cancergenome.nih.gov.

We downloaded data from TCGA using tools provided by Genomic Data Commons (GDC) [45]. We downloaded *Gene Expression Quantification* data type in *transcriptome profiling* category. We choose *RNA - Seq* with *HTSeq - FPKM* as workflow type. We downloaded the 1222 samples from TCGA-BRCA project and 1145 from TCGA-LUSC and TCGA-LUAD projects.

During the analyses of lung cancer we considered only the 408 samples with a subtype annotation available in the TCGABiolinks GUI [46].

### TCGABiolinks and metadata

We downloaded metadata for TCGA’s samples using the TCGABiolinks GUI at *Version:2*.*17*.*1* [46,47].

During the analyses of breast cancer we downloaded both the *SubtypePam50* classification labels provided by [29] and available trough `TCGAquery_subtype` function of TCGABiolinks, and *SubtypeSelected* obtained via the `PanCancerAtlas_subtypes` function of TCGABiolinks [28]. TCGABiolinks gives access to a curated table retrieved from synapse and adds some of the subtypes defined by more recent report like [48]. We discussed the performance of hSBM in classifying the two.

### Unified dataset

We downloaded data for lung from a dataset prepared by Wang et al. [38]. They processed data from GTEx and TCGA with the same pipeline and successfully corrected for study-specific biases, enabling comparative analysis. We downloaded the second version of their normalised data from figshare [40].

Only samples with a valid annotation were considered and this left us with 1415 samples from LUAD, LUSC and GTEx. We applied a log_2_(*FPKM* + 1) transformation, this reduced the number of edges *E* and let the algorithm to be faster even with a large number of nodes *N*.

In our repository we provide the code to correctly preprocess the data in order to be loaded by the model.

### Gene selection

As mentioned above we filtered genes with two different strategies.

- *Tissue specific genes* We searched genes whose behaviour was different in one tissue with respect to all the others. We used the following procedure. Firstly we estimated the mean expression of gene *g* in tissue *T* (e.g. breast)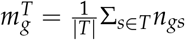, being *s* the samples and *n*_*gs*_ the expression value, the mean in the others (e.g non-breast) is 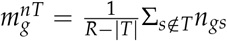 being *R* the total number of samples. Similarly, we estimated the variance of breast samples 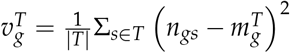. We defined the distance between the means of this two distributions as 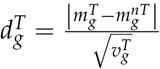. The genes with highest *d*_*g*_ were selected. Moreover, we considered only the genes that satisfied quality filters applied during eQTL analyses by the GTEx project and whose list is published in [7].
- *Highly Variable Genes* Another approach we considered is the standard selection for the so-called highly variable genes. We selected them using *scanpy* Python package [49] and kept the 3000 most variable genes.

We report in the Supplementary Figure S4 the results obtained with both strategies in each setting. We run hSBM multiple times (changing the random seed) and noticed that the peak of NMI score is quite stable and comparable with the number of classes we expected. In both settings the performance of hSBM are actually comparable between the two gene selections.

### Hierarchical Stochastic Block Model

We adapted hierarchical Stochastic Block Model (hSBM) to gene expression data. The original code to run hierarchical Stochastic Block Model on a bipartite network was provided by [9] in the repository available at: https://github.com/martingerlach/hSBM_Topicmodel/tree/develop.

Hierarchical stochastic block model is a kind of generative model that tries to maximise the probability that the model *θ* describe the data *𝒜*

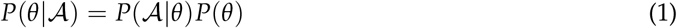

using a non-parametric approach. In the setting described in this paper 𝒜 is the gene expression matrix and the entries 𝒜_*ij*_ represents the number of FPKM of gene *i* in sample *j*. In other words 𝒜 is the adjacency matrix of a bipartite network composed by genes and samples, the edges of this network are weighted by the gene expression. The `minimise_nested_blockmodel_dl` function from the graph-tool package [50] minimises the description length Σ = *−lnP*(𝒜|*θ*) *− lnP*(*θ*) of the model. We used the nested version of the model since we expected some sort of hierarchical structure in the data [16,51,52].

We set the algorithm to minimise the description length Σ many times and selected the model that obtained the shortest description length.

As output of the model we find the probability distributions *P*(topic|sample) and *P*(gene|topic). These probabilities are defined, in terms of entries of the program as follows:

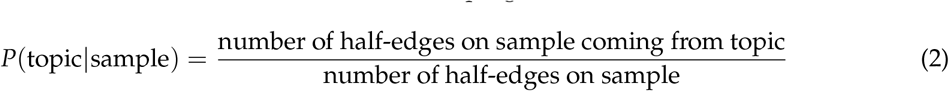

and

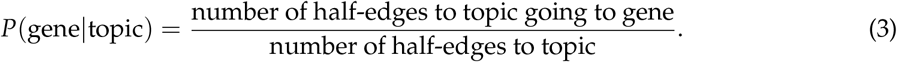

The complexity of of hSBM is *O*(*VLn*^2^*V*) if the graph is sparse, i.e. if *E ∼ O*(*V*) [16], where *V* is the number of vertices (samples and genes) and *E* the number of edges. For *E >> V* the complexity increases and the CPU time needed to minimize the description length can become prohibitive. In this case, to reduce the CPU bottleneck, one can apply a log-transformation to the data, which strongly reduces the number of edges *E*.

In our setting we have *V* = *O*(1 000) vertices, *E* = *O*(100 000) edges and the network is indeed very dense, being 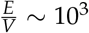. In the breast case, thanks to the strong reduction in the number of genes, we could face the task of running the program in its original setting. In the lung case, we kept 3000 genes in input and to analyze them we had to log-transform the data.

### Investigate the enrichment of the topics

We centered the *P*(topic|sample) obtaining

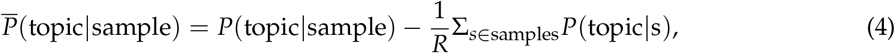

being *R* the total number of samples. This centered *P*(topic|sample) can be represented with a box plot, after grouping samples by their subtype.

A topic is nothing but a list of genes, it can be investigated using hypergeometric tests. The results shown in this paper are computed using the GSEA [34] tool.

### Survival analysis

We performed survival analysis on lung, we wanted to find how topics are related to the survival probability of a patient.

Our analysis began with the list of the mixtures *P*(topic|sample). Samples were annotated due to their subtype (LUAD or LUSC), all samples without a valid cancer type or stage label were dropped, this includes *nan* or *not reported* values. We cleaned up the labels removing any additional letter (e.g *stage ia* became *stage i*) and ended up with four stages *i, ii, iii* and *iv*.

We averaged each table over stages and obtained *P*(topic|stage) for each dataset. 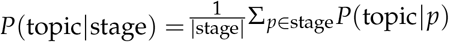 being |stage| the number of patients *p* labelled stage. We subtracted the mean to normalise the data and obtained 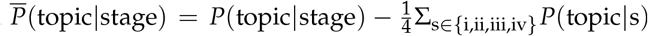. The analyses of this 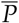 (topic|stage) was used to identify the topics with different behaviour in LUAD and LUSC.

Using GDC tools we downloaded TCGA metadata and in particular: *vital_status, days_to_last_follow_up* and *days_to_death*. We estimated the lifetime or the number of days the patient survived after the diagnosis, using *days_to_last_follow_up* if the patient was *Alive* and *days_to_death* for *Dead* patients. A similar approach was recently utilised by [41].

In order to estimate whether a topic is up regulated in a patient, we evaluated the 50th percentile of *P*(sample|topic) and considered it as a threshold *thr*. Then we engineered a feature called *up* as follows:

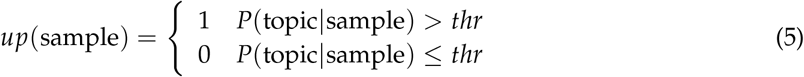

We used these data to fit the hazard with a *cox* model using the `COXPHFitter` module. We used the lifetime, the vital status and the new feature as input for the `fit` function.

The *cox* model quantified how the up regulation of the topic affected the survival probability. *Cox* fits the hazard function conditioned to a variable 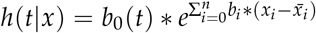 is the new feature described above. The hazard is defined as the ratio of the derivative of the survival and the survival itself 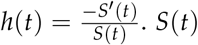 is the probability of being alive at time *t*, namely the number of patient alive at time *t* divided by the total number of patients. The package estimated the ratio between the hazard of samples with topic up-regulated and hazard of samples with topic not up-regulated. So we were able to estimate the exp(*coe f*) or hazard ratio 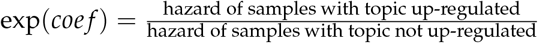. Note that the *coef* does not depend on time, but it is a sort of weighted average of period-specific hazard ratios. In our test, we obtained an exp(*coe f*) of 1.8, meaning that patients with topic up regulated have almost twice the chance to die with respect to the remaining patients. The baseline is the case in which changes of topic does not affect the survival. The P-value reported in Figure 13 is the test against this null model without other temporal dependencies. We have also checked the *gender* variable and we obtained *−* log_2_(P-value) = 1.6, which is less significant.

These analyses were performed using *lifelines* Python package[53].

### Predictor

In order to build the predictor for cancer subtypes we prepared the design matrix *X* with samples on the rows and topics on columns. The element *X*_*ij*_ is the *P*(topic_*j*_|sample_*i*_). The model was trained using Stochastic Gradient Descent so we need to apply normalisation to the matrix. We followed the standard procedure and obtained the normalised matrix 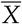 with entries 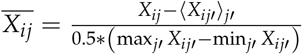 We built a one-hot encoded vector of labels corresponding to subtypes. In this analyses we dropped samples with an undefined label. We randomised and then split the data into two sets: a training and a test set. Training set was split again to obtain a validation set. In the breast setting the sizes of [train, validation, test] were in proportions of [0.18, 0.72, 0.1] and in lung [0.48, 0.32, 0.2]. The models we built in the two settings, breast and lung, are similar with some different parameters.

The model consisted of a neural network with one hidden layer. In order to keep the model as simple as possible, Stochastic Gradient Descent was selected as the optimiser in both cases. We set the loss function to be the *crossentropy*. The model was built using keras [54].

In case of breast the input layer had a dimension of 399, namely the number of topics; the hidden layer was built with 50 neurons activated by *ReLU* and the output layer consisted of 4 (one for each subtype, recall that Luminal A and B were merged into a single subtype) neurons activated by the *softmax* function.

Hyper-parameters were searched maximising the *F*1 score on the validation set. 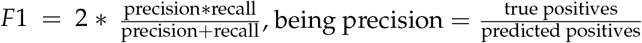, and 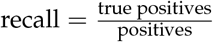.

We obtained an accuracy of 0.9008 and an Area Under Curve of 0.9798 on the test set. In the analysis of lung we prepared the design matrix *X* with the same process described above. The input layer consisted of 326 neuron, the hidden layer was instead built with 20 neurons activated by *ReLU* functions, and finally, the output layer consisted of a single neuron activated by a *sigmoid* function. The output was a binary array that distinguished LUAD and LUSC.

We obtained an accuracy of 0.9268 and an area under curve of 0.9493 on the test set.

In both cases a confusion matrix and a Receiving Operating Characteristic curve were constructed. The confusion matrix and the ROC curves were estimated using scikit learn [55]. The ROC curve represents the True Positive Rate or sensitivity 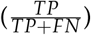 versus the False Positive Rate or 1 *–* specificity 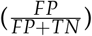 when varying the threshold on the score *Z* that defines the classes given the outputs of the hidden layer *z*. In lung *Z* is the value of the sigmoid function in the last layer *σ*(*z*). In breast we used a one-vs-all strategy and considered the softmax (*σ*_*c*_) values as probabilities for a given class *c, Z* = *σ*_*c*_(*z*).

### Implementation of WCGNA, LDA and hierarchical algorithms

In this work some of the analysis required other clustering methods. Weighted Gene Correlation Network Analysis was performed using the dedicated R package available at https://cran.r-project.org/web/packages/WGCNA/index.html. This was run using default parameters: `power` was set to the lowest for which the scale-free topology fit index curve flattens out, `minModuleSize` was set to 5 and `mergeCutHeight` to 0.2. WGCNA creates *modules* of genes, we considered them as topics. In order to obtain clusters we cut the tree built using modules to estimate distances between samples. We reported in Supplementary Figure S5 two more experiments: one when the WGCNA model is forced to behave like hierarcical clustering making small modules (low `minModuleSize`, low `mergeCutHeight`) and one where it is forced to build few big modules (high `minModuleSize`, high `mergeCutHeight`). As described in the Results section WGCNA is a valid option when the number of topics or clusters (e.g. suptypes) is well-known a-priori and, probably, a grid search of the best parameters will lead to even better results, but this is beyond the scope of this work since we wanted to compare it to hSBM which is completely non-parametric.

Latent Dirichlet Allocation (LDA) [30] is a standard and well-known topic model, we used the implementation provided by scikit-learn [56]. The model was configured using the default setting for the parameters *α* and *β*^5^: they were set to 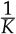, being *K* the number of topics. *K* was set based on the number of clusters in output from hSBM. When managing LDA output, we selected the *argmax* of *P*(topic|sample) to define clusters.

Hierarchical clustering was performed using *sklearn* and we set the model to use `euclidean` metric and `complete` linkage. In this case we cut the tree to match the hSBM number of clusters. In this case it has not been possible extract any information on the genes.

### Code availability

The codes, notebooks and data to reproduce this work are available on a GitHub repository at https://github.com/fvalle1/topicTCGA.

## 5. Conclusions

In conclusion, there are three properties of the hSBM topic modelling analysis that we think are at the basis of the effectiveness of this approach and correspondingly there are three lessons that we can learn from our work.

- hSBM imposes a minimal amount of assumptions on the nature of the statistical distributions of gene expression values across samples [9]. The absence of strong priors can be an important feature in clustering or topic modelling algorithms that have to be applied to complex systems. In fact, complex systems are often characterized by power-law probability distributions of frequencies [57–59] and the same is true for gene expression data [60]. Different priors, such as the ones of LDA, can drive the algorithm far from the actual system statistical properties [9]. Leveraging on the absence of simplifying priors, hSBM reaches good performances in this context and can, for example, outperform standard algorithms like LDA or WGCNA in clustering samples even if at the cost of a longer computational time.
- Cell phenotypes are driven by the expression pattern of the whole set of genes and not by a handful of markers. This is probably true in general, but it is even more plausible for complex diseases like cancer. The breast subtype classification is based on the expression levels of a handful of genes but, notwithstanding this, hSBM was able to reproduce the classification to a large extent without resorting to marker genes. Clearly this is telling us that breast cancer subtypes are driven not only by the expression level of one or two genes but by a whole pattern of pathway alterations as highlighted by the topic distribution in the different clusters. This can have important consequences on the way we approach the search for gene signatures. While it is certainly true that gene signatures play an important role in cancer medicine, it is also clear that they are not the end of the story. It is not strange to find different gene signatures aimed to identify the same cancer subtype that have almost no overlap (we have seen a prototypical example in the lung survival analysis above). This supports the importance of data mining tools able to address the overall behaviour of the transcriptome and not driven by the gene expression pattern of a single gene signature.
- Looking at all our findings we see that the actual information on the complex disease is well encoded in the topic space and in particular in the probability distributions *P*(topic|sample) and *P*(gene|topic). This is the probably the most important lesson of our analysis. An effective way to capture the complexity of a disease like cancer is through a stochastic approach in which the relationships between samples and genes are of probabilistic nature. In hSBM these complex associations are mediated by the intermediate layer of topics. This is a possible solution but non necessarily the only one. Innovative data mining tools should keep into account this observation and allow for fuzzy and complex memberships of samples across topics and genes across clusters.

We think that these lessons could be useful beyond the specific tool (hSBM) that we discussed in this paper. They should drive the conception of a new generation of innovative and network-based data mining approaches, possibly combining the best features of the different tools that we compared in this paper, such as LDA, WGCNA and hierarchical clustering. The development of new statistical tools is a mandatory task if one wants to address the challenging issue of combining the different layers of information which are available in complex databases like TCGA, say, the microRNA expression levels, the mutational content of genes or the epigenetic layer of regulation. A fruitful combination of these different sources of information could greatly enhance our ability to address a personalized therapeutic approach to complex diseases like cancer.

## Author Contributions

conceptualization, F.V., M.O. and M.C.; methodology, F.V., M.O. and M.C.; software, F.V.; writing–original draft preparation, F.V., M.C.; writing–review and editing, F.V., M.O. and M.C.; visualization, F.V.

## Acknowledgments

We would like to acknowledge the Competence Centre for Scientific Computing C^3^S which provided us the access to the computing cluster OCCAM. The results shown here are in part based upon data generated by the TCGA Research Network: https://www.cancer.gov/tcga.

## Abbreviations

The following abbreviations are used in this manuscript:
hSBM: hierarchical stochastic block model
TP, FP, TN, FN: True Positives, False Positives, True Negatives, False Negatives
FDR: False Discovery Rate
FPKM: Fragments Per Kilobase of transcript per Million mapped reads
GSEA: Gene Set Enrichment Analysis

## Footnotes

1 The Human Epidermal growth factor Receptor 2 (HER2) is a growth-promoting protein and plays an important role in several signaling pathways.

2 Ki-67 is a nuclear antigen expressed by all proliferating cells during late G1 through the M phases of the cell cycle, peaking in the G2-M and with a rapid decline after mitosis and is thus an indicator of cancer cells growth.

3 The algorithm identifies then two further levels of partition with 149 and 1204 clusters and 147 and 399 topics respectively. These partitions on the cluster side convey little information and their score is very low. They correspond to the rightmost points in the Figures 2b and c. The lowest partition level on the topic side will instead play an important role in the following when we shall discuss a classifier for breast cancer samples.

4 Cluster 6, which is also depleted, does not allow a statistical analysis since it contains too few samples.

5 *α* and *β* represent the parameters of the Dirichlet distribution from which the words (of a topic) and the topics (of a document) are sampled.

