## Supplementary material for "A topic modelling analysis of TCGA breast and lung cancer transcriptomic data"

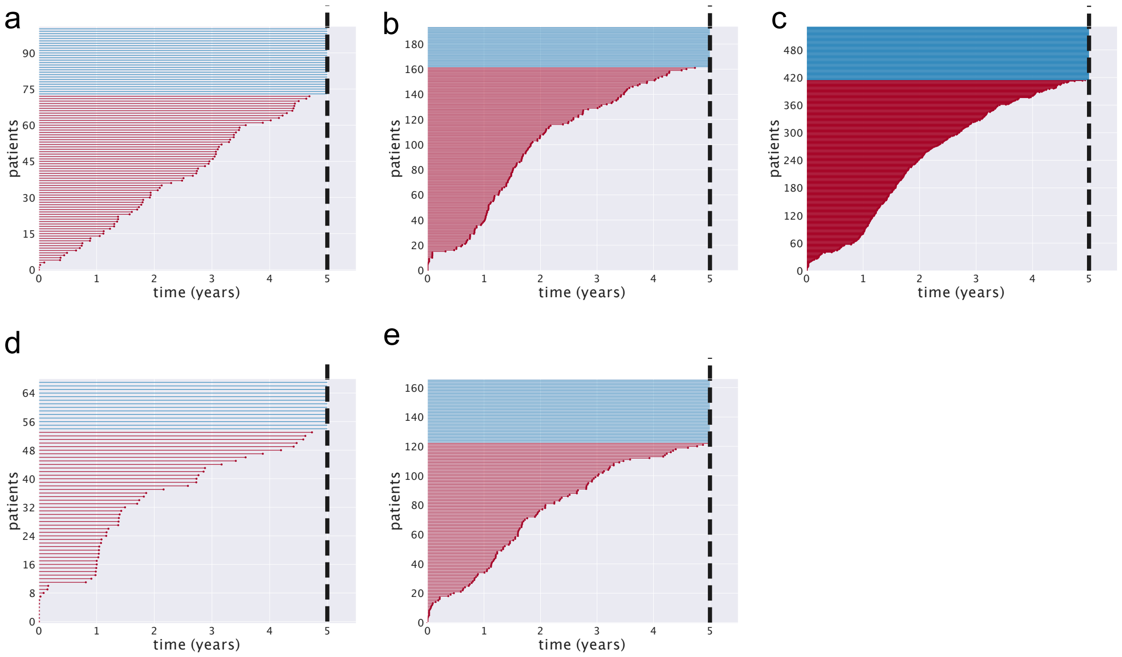


Figure S1: **Survival analysis for Breast Cancer.** We report the lifetime (number of years after diagnosis) for each sample (every horizontal line represents a sample). In blue are reported the patients that survived more than 5 years. We estimated the survival probability as the fraction of patients alive after 5 years. We compared this value with a null model obtained by reshuffling the labels multiple times and evaluated a Z-score in this way. We obtained a survival of: 0.28 ($Z = 1.93$) for Normal-like in (**a**), 0.16 ($Z = 2.03$) for Luminal B in (**b**), 0.22 ($Z = 0.25$) for Luminal A in (**c**), 0.21 ($Z = 0.26$) for HER2 in (**d**) and 0.26 ($Z = 1.54$) for Basal in (**e**).

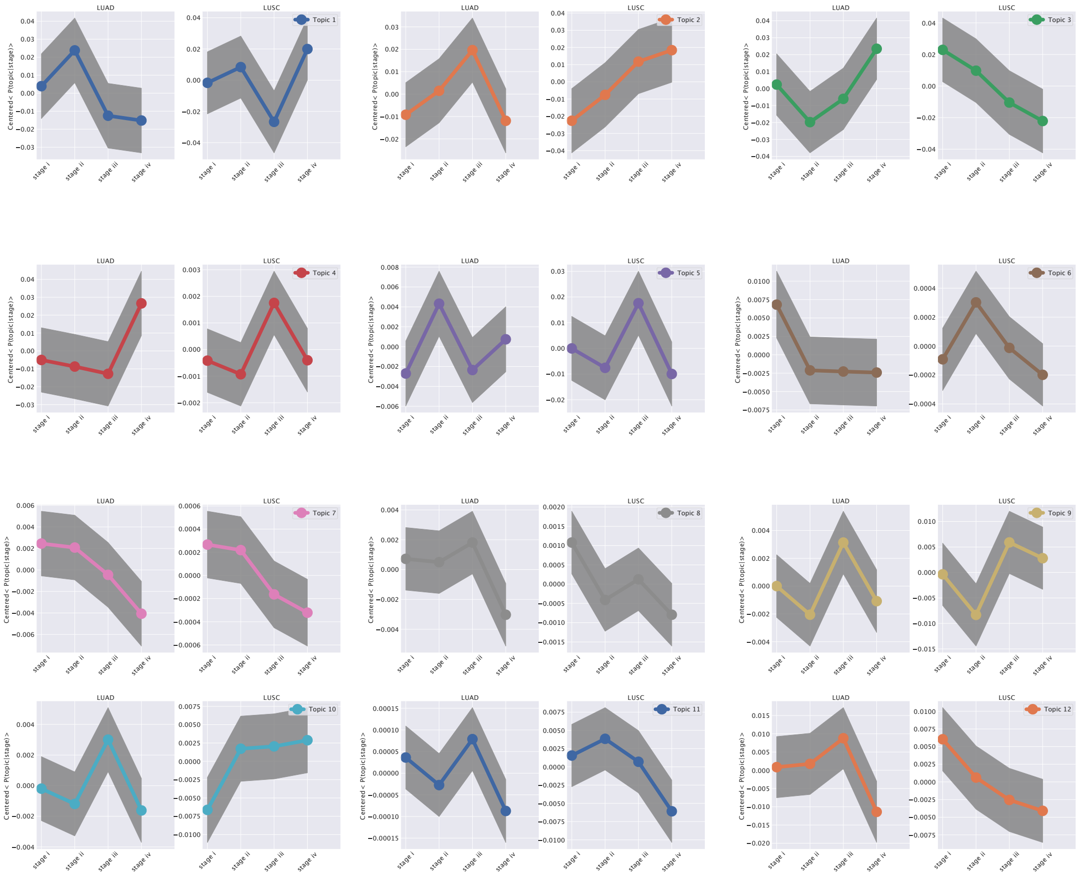


Figure S2: **Survival analysis for Lung cancer.** We analysed the trend of $P(topic|stage)$ in LUAD and LUSC patients. We searched topics in which the trend showed a difference in the two cohorts. We identified *Topic 3* as our candidate topic, since it is the one in which the importance of the topic increases in LUAD and decreases in LUSC.

| | PAM50 | Subtype Selected |
|---|---|---|
| Basal | 212 | 188 |
| HER2 | 91 | 82 |
| Luminal A | 633 | 576 |
| Luminal B | 231 | 217 |
| Normal-like | 42 | 142 |

Figure S3: **Number of samples per each annotation in Breast cancer.** TCGABiolinks assigns more Normal-like subtypes.

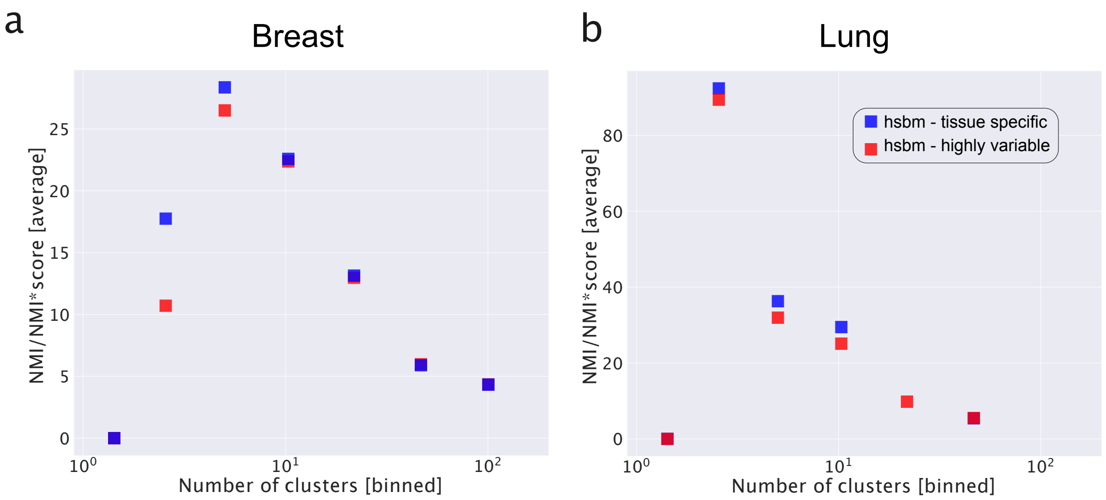


Figure S4: **Performance of hSBM with different gene selections.** We ran hSBM 10 times and looked at the average score obtained at different resolutions. We estimated the average score at any number of clusters (we made uniform bins and averaged all the scores obtained at such resolution). The figures report the comparison made running hSBM with tissue specific genes or highly variable genes as discussed in the methods section of the main text. In (**a**) the results for Breast and in (**b**) for Lung cancer. The two selections are almost compatible being the tissue specific a bit more performing. Moreover it is interesting to notice that the highest score was obtained for a number of cluster similar to the number of subtypes (5 for Breast and 2 for Lung). Note that the runs are not all equivalent, the description length of the model can be different. The results reported in the main text refer to the runs with the shortest description length (i.e. the models which need the least number of bits to describe the data).

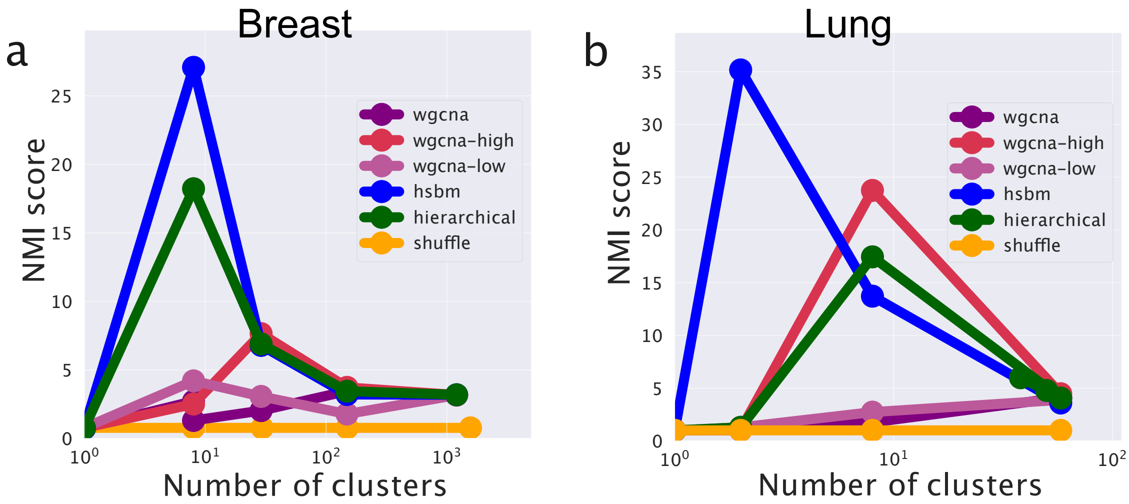


Figure S5: **Performance of WGCNA with different settings.** We ran WGCNA with different thresholds. `wgcna-high` represents a configuration in which WGCNA is set to find an high number of modules (topics): this case it similar to hierarchical clustering. `wgcna-low` represents a setting in which the algorithm is set to found few modules. The label `wgcna` represents the setting reported in the paper in which we set it to emulate the number of topics of hSBM (which searches the optimal number of topics itself). When WGCNA searches many modules, its outcome is similar to the hSBM (and of course to hierarchical clustering) one. Therefore, if WGCNA is set to replicate the resolution (i.e., the number of topics and clusters) of hSBM its classification performances are low.
